# Shared risk alleles with discordant polygenic effects: Disentangling the genetic overlap between ASD and ADHD

**DOI:** 10.1101/580365

**Authors:** Ellen Verhoef, Jakob Grove, Chin Yang Shapland, Ditte Demontis, Stephen Burgess, Dheeraj Rai, Anders D. Børglum, Beate St Pourcain

## Abstract

Insight into shared polygenetic architectures affects our understanding of neurodevelopmental disorders. Here, we investigate evidence for pleiotropic mechanisms that may explain the comorbidity between Autism Spectrum Disorder (ASD) and Attention-Deficit/Hyperactivity Disorder (ADHD). These complex neurodevelopmental conditions often co-occur, but differ in their polygenetic association patterns, especially with educational attainment (EA), showing discordant association effects. Using multivariable regression analyses and existing genome-wide summary statistics based on 10,610 to 766,345 individuals, we demonstrate that EA-related polygenic variation is shared between ASD and ADHD. We show that different combinations of the same ASD and ADHD risk-increasing alleles can simultaneously re-capture known ASD-related positive and ADHD-related negative associations with EA. Such patterns, although to a lesser degree, were also present for combinations of other psychiatric disorders. These findings suggest pleiotropic mechanisms, where the same polygenic sites can encode multiple independent, even discordant, association patterns without involving distinct loci, and have implications for cross-disorder investigations.

## Main

Autism Spectrum Disorder (ASD) and Attention-Deficit/Hyperactivity Disorder (ADHD) are genetically complex childhood-onset neurodevelopmental disorders^1,2^ that often co-occur^3^. Approximately 15–25% of individuals with ADHD show ASD symptoms, and ~40–70% of individuals with ASD have a comorbid ADHD symptomatology^3^.

Both rare and common genetic variation contributes to ASD and ADHD liability^4–7^. There is increasing evidence from twin and molecular studies^8,9^ suggesting genetic links between ASD and ADHD symptoms, both throughout population variation^10–16^ and at the extremes^10,17^. The existence of genetic cross-disorder links is further strengthened by the familial co-aggregation of both clinical disorders in large register-based studies^18^. Consistently, the identification of shared copy number variations (CNVs) in ASD and ADHD suggests similar biological pathways^19^. Estimates of cross-disorder genetic correlations range between 0.54 and 0.87 in twin analyses^20^. When studying polygenic variation, symptom overlap can even be stronger in population-based samples^11^, although these links are notably lower between clinically defined ASD and ADHD^21–23^. While recent research has reported moderate genetic correlations between clinical ASD and ADHD, based on genome-wide summary statistics (r_g_=0.36)^21^, earlier studies with lower statistical power found little evidence for such genetic overlap^22,23^.

Besides some shared genetic aetiology, there are differences in the polygenic architecture of clinical ASD and clinical ADHD. Each disorder, as tagged by common variants, shows an opposite genetic correlation with cognitive functioning and educational attainment (EA), where the latter is influenced by both cognitive abilities and socio-economic status (SES)^24^. While increased polygenic ADHD risk has been linked to lower cognitive abilities and EA^25–28^, increased polygenic ASD risk has been associated with higher cognitive functionality and EA^21,27,29^. This discordant association pattern is strongest for measures of years-of-schooling and college-completion^21,28^. Recent evidence for a polygenic p-factor, shared across major psychiatric disorders including ASD and ADHD^30^, suggests overarching genetic similarities between neurodevelopmental disorders. This may also involve shared polygenic variation among ASD, ADHD and EA, acting through complex pleiotropic, mediating or confounding mechanisms. However, the discordant association profile of each disorder with EA might also be explained by independent genetic loci.

To improve our understanding of shared psychopathologies across disorders, this study aims to disentangle the genetic overlap between ASD and ADHD with respect to EA-related associations using a multivariable regression (MVR) framework and individual (not accumulated^31^) SNP-based information from existing genome-wide association study (GWAS) summary statistics. This translates a causal modelling approach^32^ into a polygenic context without making causal inferences. We dissect polygenic associations between each disorder and EA into either ASD-specific or ADHD-specific associations as well as genetic influences that are shared across both disorders and EA. We assess the specificity of these association profiles by examining combinations of other psychiatric disorders and finally model their impact on cross-disorder investigations.

## Results

### Multivariable regression model fitting

Modelling the effect of ASD-related risk alleles on EA (ASD-MVR), we jointly estimated ASD-specific associations as well as cross-disorder associations shared with ADHD (Figure 1a). Similarly, modelling the effect of ADHD-related risk alleles on EA (ADHD-MVR), we jointly assessed ADHD-specific associations as well as cross-disorder associations shared with ASD (Figure 1b). Compared to univariable ASD and ADHD regression models (modelling polygenic associations between a single disorder and EA only), ASD-MVRs and ADHD-MVRs allowing for cross-disorder associations with EA revealed a better model fit. They explained up to 3% more variation in genetically predictable EA, with little evidence for multi-collinearity (Supplementary Table 8-10).

**Figure 1:**
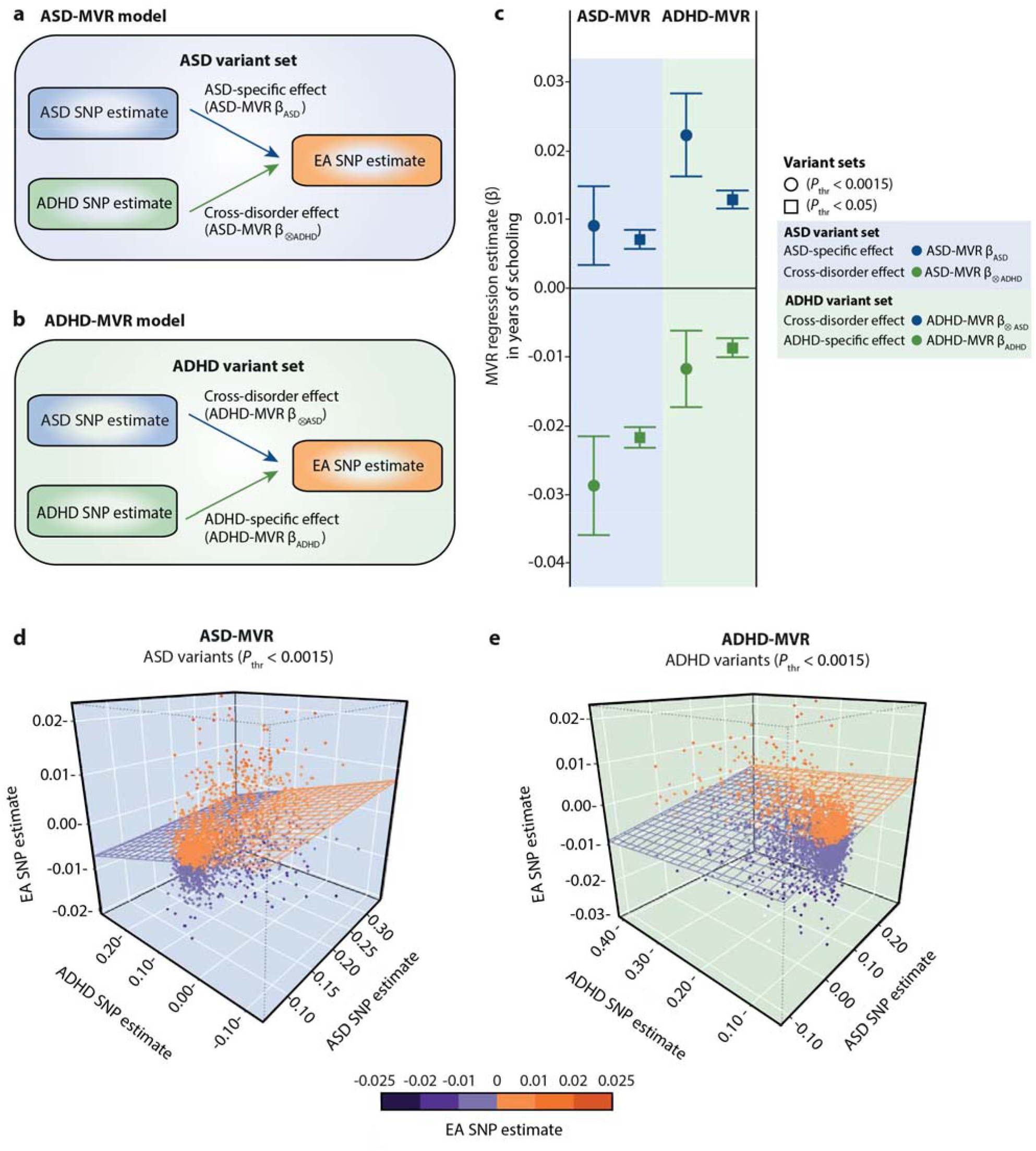
ASD-specific, ADHD-specific and cross-disorder associations with educational attainment. Sets of independent ASD and ADHD genetic variants were selected from ASD(iPSYCH, woADHD) and ADHD(iPSYCH) GWAS statistics respectively, across different *P*-value thresholds (*P*_thr_<0.0015, *P*_thr_<0.05). Corresponding SNP estimates for ASD, ADHD and EA were subsequently extracted from ASD(iPSYCH, woADHD), ADHD(iPSYCH) and EA(SSGAC) GWAS statistics respectively. **(a)** Schematic ASD-MVR model jointly estimating ASD-specific (ASD-MVR β_ASD_) and ADHD cross-disorder (ASD-MVR β_⊗ADHD_) associations with EA using ASD variant sets. ASD-MVR β_ASD_ were fitted with ASD SNP estimates, ADHD cross-disorder effects (ASD-MVR β_⊗ADHD_) were fitted with ADHD SNP estimates. For clarity, intercepts are not shown. **(b)** Schematic ADHD-MVR model jointly estimating ADHD-specific (ADHD-MVR β_ADHD_) and ASD cross-disorder (ADHD-MVR β_⊗ASD_) associations with EA using ADHD variant sets. ADHD-MVR β_ADHD_ were fitted with ADHD SNP estimates. ASD cross-disorder effects (ADHD-MVR β_⊗ASD_) were fitted with ASD SNP estimates. For clarity, intercepts are not shown. **(c)** Estimated ASD-specific (ASD-MVR β_ASD_), ADHD-specific (ADHD-MVR β_ADHD_) and cross-disorder association effects (ASD-MVR β_⊗ADHD_, ADHD-MVR β_⊗ASD_) with SNP estimates aligned according to ASD risk (ASD-MVR, 1a) and ADHD risk (ADHD-MVR, 1b), respectively. All MVR effects are presented as change in years-of-schooling per increase in log-odds ASD or ADHD liability respectively. Bars represent 95% confidence intervals. **(d)** 3D scatter plot of ASD SNP estimates (lnOR, x-axis), ADHD SNP estimates (lnOR, y-axis) and EA SNP estimates (z-axis) for ASD-related variants (*P*_thr_<0.0015). The multivariable regression plane reflects ASD-specific and ADHD cross-disorder associations, as shown in 1c. **(e)** 3D scatter plot of ASD SNP estimates (lnOR, x-axis), ADHD SNP estimates (lnOR, y-axis) and EA SNP estimates (z-axis) for ADHD-related variants (*P*_thr_<0.0015). The multivariable regression plane reflects ADHD-specific and ASD cross-disorder associations, as shown in 1c. All presented MVR effects passed the multiple testing threshold of *P*<0.0023. Abbreviations: ADHD, Attention-Deficit/Hyperactivity Disorder; ASD, Autism Spectrum Disorder; EA, educational attainment; MVR, multivariable regression; *P*_thr_, *P*-value threshold.

### Multivariable regression analyses of EA on ASD and ADHD

Discovery ASD-MVRs and ADHD-MVRs were carried out with a series of variant sets covering different *P*-value selection thresholds (11 variant sets for each disorder: 5×10^−8^<*P*_thr_<0.5, Supplementary Figure 1a), similar to a polygenic scoring approach^31^, and provided evidence for ASD-specific, ADHD-specific and cross-disorder associations with EA (Supplementary Table 6-7). For example, for ASD-MVR at *P*_thr_<0.0015 (N_SNPs_=1,973, Figure 1a,c, Supplementary Table 8), we observed an 0.009 increase in years-of-schooling per log-odds in ASD-liability (ASD-MVR β_ASD_=0.009(SE=0.003), *P*=0.002), and a 0.029 decrease in years-of-schooling per log-odds in ADHD-liability (ASD-MVR β_⊗ADHD_=−0.029(SE=0.004), *P*<1×10^−10^). Thus, these cross-disorder associations showed an opposite direction of effect (Figure 1c,d), even though they were modeled with the same ASD-related risk alleles.

An analogous approach with ADHD-MVRs (Figure 1b) revealed a complementary association profile (Supplementary Table 7). There was an inverse ADHD-specific association between polygenic ADHD risk and EA. Conditionally, ASD cross-disorder associations with EA were positive, thus discordant, even though modeled with the same ADHD-related risk alleles (Figure 1c,e). For ADHD-MVR at *P*_thr_<0.0015 (N_SNPs_=2,717, Figure 1c,e, Supplementary Table 8), this corresponds to an 0.012 decrease in years-of-education per log-odds in ADHD liability (ADHD-MVR β_ADHD_=-0.012(SE=0.003), *P*=4×10^−5^), and an increase in 0.022 years-of-education per log-odds in ASD liability (ADHD-MVR β_⊗ASD_=0.022(SE=0.003), *P*<1×10^−10^). Importantly, joint modelling of disorder-specific and cross-disorder SNP effects as part of a multivariable regression model also increased the evidence for ASD-specific and ADHD-specific association effects (Supplementary Table 8). While disorder-specific and cross-disorder MVR effects are independent of each other, the underlying genetic variation is shared and the simultaneous estimation of ASD and ADHD MVR effects results in an improved model fit. Increasing the number of variants in ASD-MVRs and ADHD-MVRs using more relaxed SNP-selection thresholds (e.g. *P*_thr_<0.05) boosted the statistical power (Figure 1c, Supplementary Table 6-8).

Follow-up analyses with concordant variants (variants with concordant ASD and ADHD risk effects, ~80% of the initial sets) confirmed these findings (Supplementary Figure 1b, Supplementary Table 9) and showed that MVR findings are independent of allelic alignment to ASD or ADHD risk. Bivariate relationships between SNP estimates for ASD, ADHD and EA are plotted in Supplementary Figure 2 (concordant variants, *P*_thr_<0.05).

Using the same sets of variants as in the discovery ASD-MVRs and ADHD-MVRs above, we replicated the profile of discordant cross-disorder associations with EA at the relaxed threshold (*P*_thr_<0.05), using SNP estimates from ASD(PGC), instead of ASD(iPSYCH,woADHD) as predictor (Supplementary Figure 1c, Supplementary Table 10). Thus, despite known zero genetic correlations between ADHD(iPSYCH) and ASD(PGC)(Supplementary Table 3), we observed strong evidence for genetic associations (*P*<1×10^−10^) between each disorder and EA using the same set of SNPs (*P*_thr_<0.05, Supplementary Table 10). At the more stringent threshold (*P*_thr_<0.0015), only ASD-specific effects passed the multiple testing threshold. This is consistent with the limited power of ASD(PGC) and a reduced concordance rate between ASD(PGC) and ADHD(iPSYCH) risk alleles (~50%). ASD-MVRs and ADHD-MVRs including general intelligence as the outcome (Supplementary Figure 1d), instead of EA, confirmed the association patterns of the discovery MVRs throughout (Supplementary Table 11), and our findings agree with known genetic correlations (Supplementary Table 5).

### Identification of cross-disorder loci

To identify variants exerting the largest cross-disorder effects, we systematically assessed the overlap between ASD-MVR and ADHD-MVR variant sets, based on the powerful iPSYCH samples (Supplementary Figure 1e). Starting with ASD-MVR and ADHD-MVR variant sets at *P*_*th*r_<0.0015 (ASD: N_SNPs_≤1,973, ADHD: N_SNPs_≤2,717) we successively restricted the selected markers to variants that were also associated with the other disorder (5×10^−8^≤*P*_thr_<0.5, Figure 2a). Fitting MVR models with these reduced sets (Figure 2b, Supplementary Table 12-13), we identified MVR effects that were larger than in the discovery ASD-MVRs and ADHD-MVRs, with non-overlapping 95% confidence intervals. For example, for variants selected at *P*_*th*r_<0.0015 for both disorders, we estimated about 5-fold larger MVR effects, using only 4.2% and 3.1% of the original variant sets for ASD-MVR and ADHD-MVR respectively. These reduced variant sets comprised the same 83 loci, based on identical or tagged proxy SNPs (Linkage Disequilibrium-r^2^=0.6, 500 kb window), with 99% of them showing concordant ASD and ADHD risk effects (Supplementary Table 14). This combination of risk alleles and effects (selected at *P*_*th*r_<0.0015 for both disorders) is unlikely to be due to chance, as shown by permutations (Supplementary Table 15, empirical *P*<3×10^−4^), and suggests locus specificity. The 83 variants mapped to 52 genes using positional mapping and included multiple regulatory RNAs (Supplementary Table 14).

**Figure 2:**
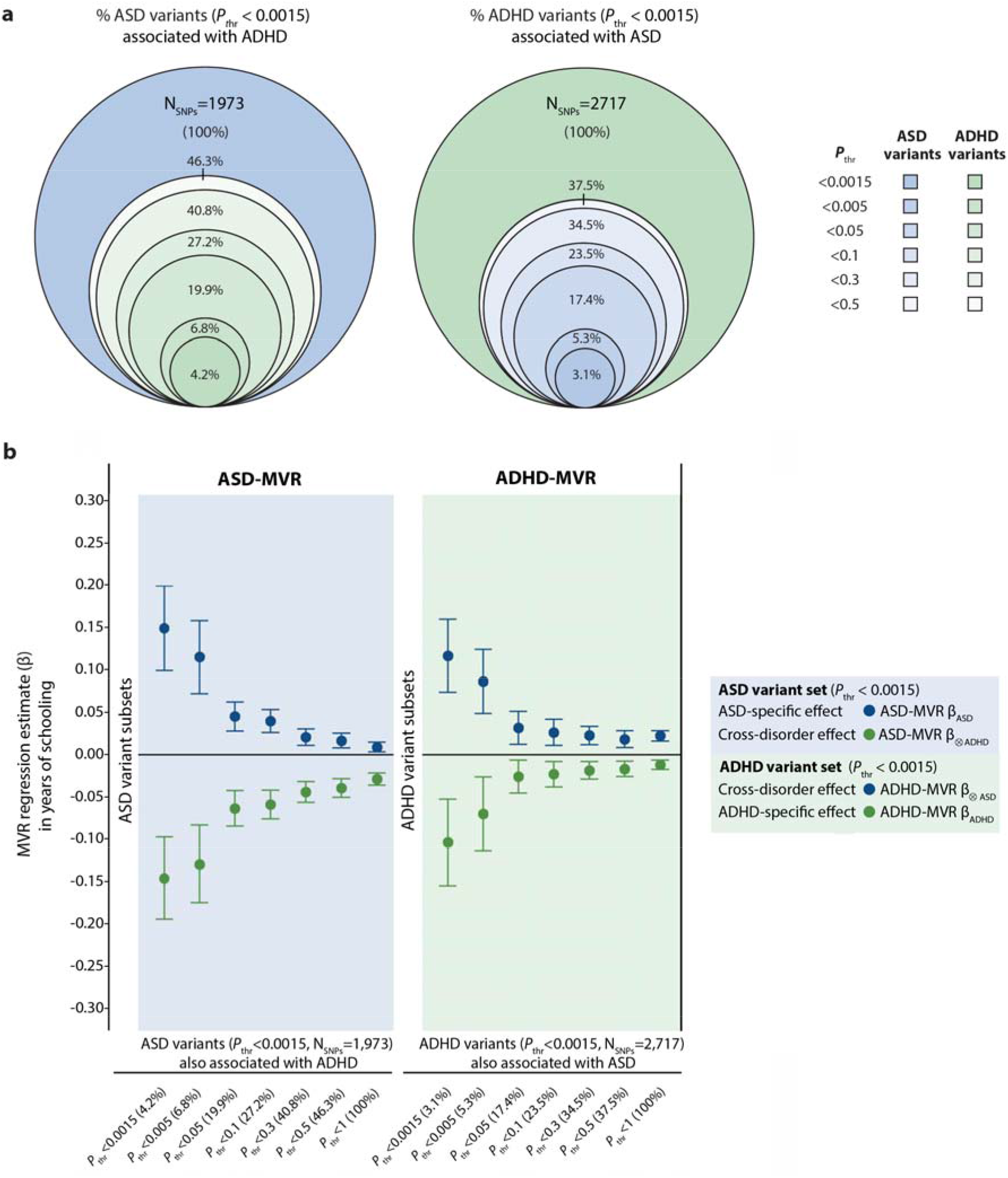
Changes in ASD-specific, ADHD-specific and cross-disorder associations with educational attainment depending on variant sets meeting joint ASD and ADHD selection criteria. (a) Percentage of ASD variants (*P*_thr_<0.0015) also associated with ADHD across multiple *P*-value selection thresholds (*P*_thr_: 0.0015; 0.005; 0.05; 0.1; 0.3; 0.5) (b) Percentage of ADHD variants (*P*_thr_<0.0015) also associated with ASD across multiple *P*-value selection thresholds (*P*_thr_: 0.0015; 0.005; 0.05; 0.1; 0.3; 0.5) (c) ASD-MVR and ADHD-MVR analyses based on SNP sets shown in (a) and (b). SNP estimates were extracted from ASD(iPSYCH, woADHD), ADHD(iPSYCH) and EA(SSGAC) GWAS statistics. Variant sets with *P*-value selection thresholds *P*_thr_ 5×10^−8^, 5×10^−7^, 5×10^−6^, 5×10^−5^ and 0.0005 for the cross-disorder are not shown, due to the small number of variants identified. All MVR effects passed the multiple testing threshold of *P*<0.0011, except ADHD-specific effects for ADHD-MVR (where variants were associated with ASD at *P*_thr_: 0.005, 0.05, 0.1). Abbreviations: ADHD, Attention-Deficit/Hyperactivity Disorder; ASD, Autism Spectrum Disorder; MVR, multivariable regression; *P*_thr_, *P*-value threshold.

### Specificity of ADHD/ASD cross-disorder genetic associations

To assess the specificity of discordant ADHD/ASD cross-disorder associations with EA, based on the selected shared risk allele pool, we also modelled cross-disorder associations between other neuropsychiatric disorders (MDD, SCZ and BD) and EA, using the previously defined ASD and ADHD variant sets (*P*_thr_<0.0015 and *P*_thr_<0.05, Supplementary Figure 1f). This identified several similar association patterns, predominantly at *P*_thr_<0.05 (Supplementary Figure 3, Supplementary Table 16-17), each consistent with known genetic correlations (Supplementary Table 3-4). For ASD-MVRs at *P*_thr_<0.05, discordant patterns with EA were detected in combination with MDD (ASD-MVR β_⊗MDD_=-0.012, SE=0.001, *P*<1×10^−10^, Supplementary Table 16). For ADHD-MVRs at *P*_thr_<0.05, discordant associations with EA were found with respect to BD (ADHD-MVR β_⊗BD_=0.008, SE=0.001, *P*<1×10^−10^, Supplementary Table 17). Identified cross-disorder effects were smaller compared to ASD/ADHD cross-disorder associations with EA, observed in the discovery MVRs based on iPSYCH (Figure 1c, Supplementary Figure 3), but comparable to follow-up analyses using ASD (PGC).

### Multi-factor model of genetic interrelations between ASD, ADHD and educational attainment

Consistent with an assumption-free decomposition of trait-interrelationships (Cholesky model), our findings support a multi-factor model that predicts at least two sources of shared genetic variation between ASD and ADHD (Figure 3). The first genetic factor (A1, EA/ADHD/ASD) captures shared variation between EA, ASD and ADHD and predicts negative genetic covariance between ASD and ADHD, consistent with MVR findings in this study. The second genetic factor (A2, ADHD/ASD) acts independently of A1 and explains positive genetic covariance between ASD and ADHD, reflecting known positive or null genetic correlations between disorders^21,22^. The third genetic factor (A3) encodes ASD-specific variation. The observed net covariance between ASD and ADHD reflects the sum of negative and positive covariance contributions. Consequently, ASD/ADHD genetic overlap might be reduced, as hypothesised for ASD(iPSYCH)/ADHD(iPSYCH)(Figure 3a). It might also be completely abolished, as hypothesised for ASD(PGC)/ADHD(iPSYCH)(Figure 3b) and supported by simulations that recapture genetic trait interrelationships (Supplementary Table 18). Alternative definitions of the model can allow for ADHD-specific influences (Supplementary Figure 5).

**Figure 3:**
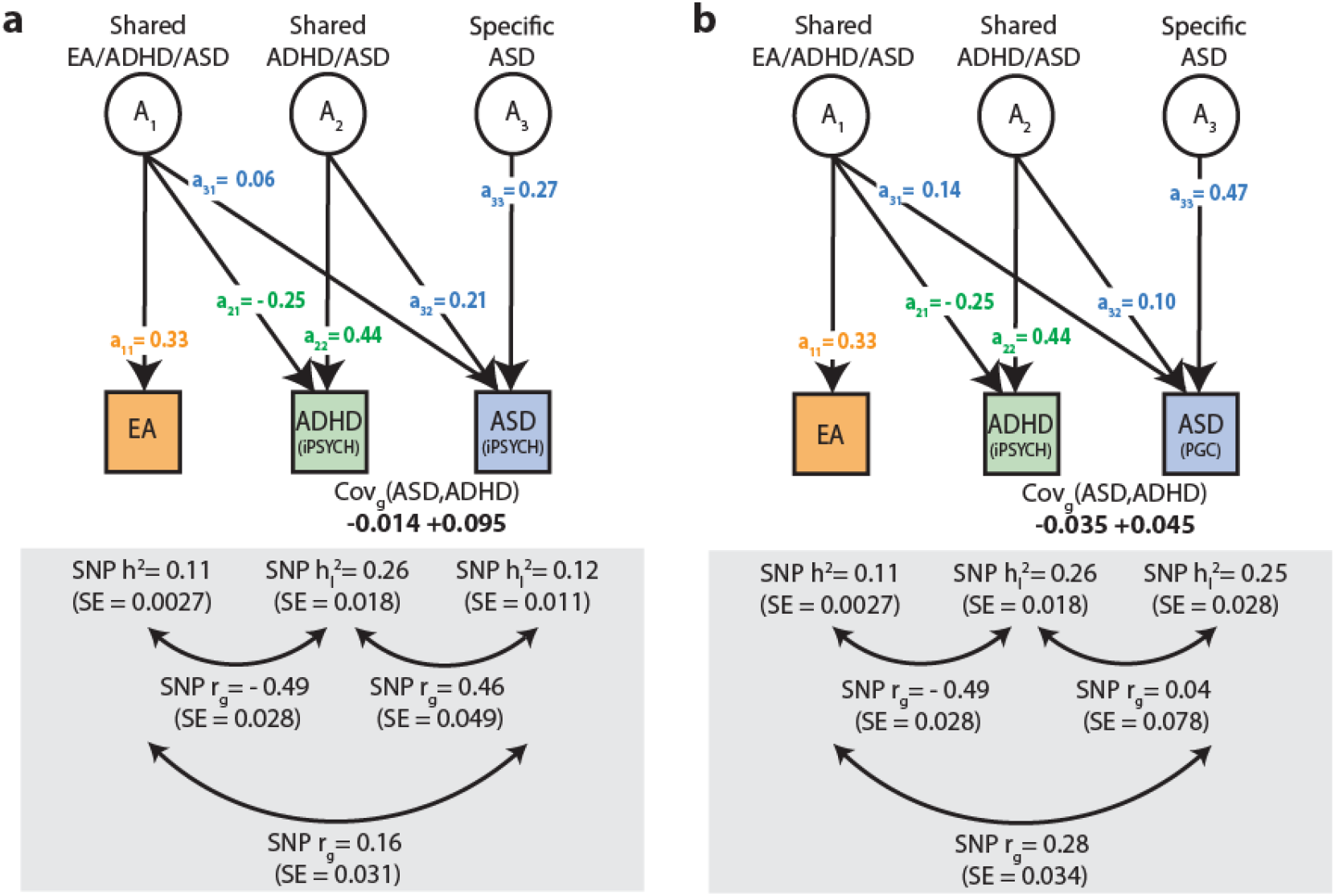
Multi-factor model of genetic interrelations between ASD, ADHD and educational attainment. The model predicts two sources of shared genetic influences between ASD and ADHD, as captured by common variants within an infinitely large population. The first genetic factor (A1, shared EA/ADHD/ASD) refers to shared genetic variation between EA, ADHD and ASD. It allows for a negative genetic covariance between ASD and ADHD. The second genetic factor (A2, shared ADHD/ASD) acts independently of A1, explaining positive genetic covariance between ASD and ADHD. Each factor loading (“a”) for the Cholesky decomposition of a trivariate trait is described in the Methods. **(a)** Multi-factor model consistent with ASD(iPSYCH), ADHD(iPSYCH) and EA(SSGAC) summary statistics. **(b)** Multi-factor model consistent with ASD(PGC), ADHD(iPSYCH) and EA(SSGAC) summary statistics and supported by simulations (Supplementary Table 18). Factor loadings (“a”) were derived from LDSC SNP-heritabilities and genetic correlations. Shared ADHD/ASD genetic influences (A2) were modelled allowing for ASD-specific effects (A3). Phenotypic measures are represented by squares, while latent genetic factors are represented by circles. Single-headed arrows denote genetic factor loadings (“a”), double-headed arrows genetic correlations (“r_g_”). Residual influences and unit variances for latent variables were omitted. Abbreviations: ADHD, Attention-Deficit/Hyperactivity Disorder; ASD, Autism Spectrum Disorder; EA, educational attainment; iPSYCH, The Lundbeck Foundation Initiative for Integrative Psychiatric Research; PGC, Psychiatric Genomics Consortium; SNP h^2^, SNP heritability; SNP r_g_, SNP genetic correlation, cov_g_, genetic covariance; woADHD, without ADHD

The predicted multi-factor model is furthermore consistent with genetic correlations for ASD+ADHD symptom combinations. For ASD(iPSYCH,woADHD), excluding comorbid ADHD patients, genetic correlations with EA exceeded those between ASD(iPSYCH) and EA, although the 95%-confidence intervals overlap (Figure 4). In contrast, inverse genetic correlations between ADHD and EA (r_g_=−0.49(SE=0.03), *P*<1×^−10^) were dampened when ADHD(iPSYCH) and ASD(PGC) summary statistics were combined and then correlated with EA (r_g_=−0.33(SE=0.03), *P*<1×10^−10^)(Figure 4).

**Figure 4:**
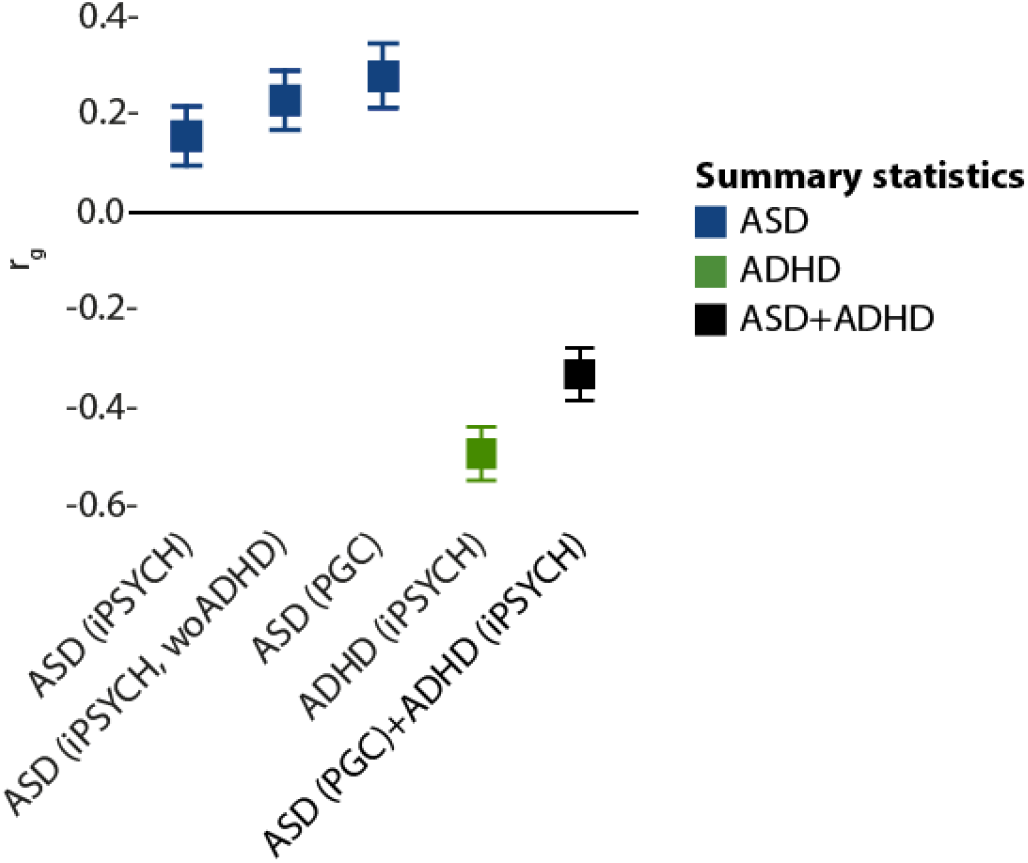
Genetic correlations between educational attainment, ASD and ADHD. Genetic correlations of ASD and ADHD samples with educational attainment (EA) were estimated using genome-wide summary statistics for EA(SSGAC), ASD(iPSYCH), ASD(iPSYCH, woADHD), ASD(PGC), ADHD(iPSYCH) and ASD(PGC)+ADHD(iPSYCH) respectively. Latter were created by performing a random-effect meta-analysis combining ASD(PGC) and ADHD(iPSYCH) genome-wide summary statistics. Genetic correlations were estimated with unconstrained LD score correlation. Bars represent 95% confidence intervals. Abbreviations: ASD, Autism Spectrum Disorder; ADHD, Attention-Deficit/Hyperactivity Disorder; ASD (iPSYCH,woADHD), ASD without ADHD; iPSYCH, The Lundbeck Foundation Initiative for Integrative Psychiatric Research; PGC, Psychiatric Genomics Consortium

## Discussion

This study provides strong and consistent evidence that EA-related polygenic variation is shared across ASD and ADHD. Different combinations of the same risk-increasing alleles can result in ASD-related positive and ADHD-related negative association profiles with genetically predictable EA. This suggests the presence of pleiotropic mechanisms, where multiple, even discordant, association profiles with EA can be encoded across the same polygenic sites without involving distinct loci.

The pattern of ASD- and ADHD-specific associations with EA, in combination with discordant polygenic cross-disorder links, was (i) reciprocally detectable using both ASD and ADHD-related variant sets, (ii) replicated at *P*_thr_<0.05 using ASD(PGC) summary statistics, (iii) independent of risk allele alignment for ASD and ADHD and (iv) consistent with the previously reported genetic overlap between EA, ASD and ADHD^21,28^. This suggests that cross-disorder associations are driven by a substantial proportion of subthreshold variants that are associated with both ASD and ADHD, reflecting pleiotropic effects, presumed for many trait-associated variants in the genome^33^. Moreover, joint modelling of these pleiotropic alleles, exploiting single instead of aggregated SNP estimates, could substantially increase evidence for both disorder-specific and cross-disorder associations. Against this shared polygenic background involving several thousands of variants, we also identified ~80 loci (~50 genes) that exerted discernably larger signals when followed-up in the powerful iPSYCH samples, and involve regulatory loci.

The identification of discordant ASD- and ADHD-related association profiles with EA, across shared ADHD and ASD risk alleles, may relate to different mechanisms. First, there is mounting evidence that ASD and ADHD share some underlying aetiological mechanisms. For example, CNVs in ASD and ADHD indicate similar biological pathways^19^, and both disorders carry a similar burden of rare protein-truncating variants, implicating shared genes^34^. Despite these biological commonalities, the assignment of clinical diagnoses to patients comorbid for ASD and ADHD symptomatology has been, until the introduction of the Diagnostic Statistical Manual of Mental Disorders 5th edition (DSM-5)^35^, less formalised. GWAS participants have been predominantly diagnosed with previous classification systems^36,37^, where hierarchical diagnostic criteria did not allow for a diagnosis of ADHD when symptoms occurred during the course of a pervasive developmental disorder. Furthermore, comorbid symptoms within clinical ASD and ADHD often occur at the subthreshold level^3^. This suggests that patients with comorbid ASD and ADHD symptoms might have been assigned to either diagnostic entity, depending on the symptoms that presented first, potentially exacerbating genetic similarities between ASD and ADHD.

Second, shared alleles with opposite polygenic effects may implicate epistasis, such that ASD-specific and ADHD-specific genetic factors may shape the direction and magnitude of ADHD/ASD cross-disorder associations with EA. For example, following an omnigenic model^38,39^, disorder-specific ‘peripheral’ genetic influences could control shared ADHD/ASD cross-disorder ‘core’ variation. Alternatively, the set of shared risk alleles may harbour high plasticity genes, exerting different effects within differing environments. The strongest signals driving the observed opposite cross-disorder associations with EA in iPSYCH were found near several miRNA and lncRNA loci that can be influenced by environmental signals^40^. Thus, symptoms and behavioural spectrum of an individual at high genetic risk for both ASD and ADHD may depend on the exposure to different home environments (e.g. household income, neighbourhood SES). This is consistent with findings of stress-related gene modulatory effects manifesting, for example, as an environment-induced development of depression^41^.

Discordant associations with EA, encoded via different combinations of the same risk alleles, may lead to negative genetic covariance between ASD and ADHD that can reduce the net genetic overlap between both disorders. The discovered inverse cross-disorder associations with EA were stronger and larger for ADHD and ASD, when studied using iPSYCH samples, compared to cross-disorder effects involving other psychiatric disorders. However, they are unlikely to be limited to polygenic ASD and ADHD risk. The identification of discordant EA association profiles for ASD and MDD risk, across the same ASD risk alleles, and likewise the discordant polygenic EA association effects for ADHD and BD, across the same ADHD risk alleles, supports the widespread pleiotropy among neuropsychiatric disorders^30^. In particular, it suggests complex genetic interrelationships between ASD and MDD, and between ADHD and BD that may involve negative covariance.

Discordant ADHD/ASD cross-disorder association profiles with EA, across shared polygenic sites, were replicated using two independent ASD collections at a widely established selection threshold (*P*_thr_<0.05) often used in related polygenic scoring analyses^42^. This suggests that our findings are robust across diagnostic classification systems for clinical ASD, routes of patient ascertainment, and association analysis designs. Nonetheless, differences in concordance rates between ASD and ADHD risk alleles with respect to ASD(PGC)(~50%) versus ASD(iPSYCH,woADHD)(~80%) may also suggest genetic heterogeneity among ASD samples. In addition, our results could be affected by presentation bias, such that children with ASD might be more often labelled with ADHD, due to a higher proportion of ADHD symptoms in ASD^43^. Furthermore, controls are shared across iPSYCH GWAS samples, potentially leading to inflated type-I error^44,45^. This is, however, unlikely, given the opposite direction of effect and replication within ASD(PGC). Finally, as symptom heterogeneity may shape genetic overlap between neurodevelopmental disorders, EA and cognition-related traits^21,46^, future studies with access to this information are warranted to fully understand the underlying complex multivariate interrelations.

## Conclusions

We show that EA-related polygenic variation is shared across ASD and ADHD genetic architectures, and that different combinations of the same risk alleles can encode ASD-related positive and ADHD-related negative association profiles with EA without involving further loci. These inverse patterns may affect the detectable net genetic overlap between ASD and ADHD but also other psychiatric disorders, especially when patients are jointly analysed within cross-disorder investigations.

## Methods

### Data sets

Information on SNP-EA, SNP-general intelligence and SNP-disorder associations was obtained from GWAS summary statistics^21,27,28,47–49^. These aggregated results are briefly summarised here and described in detail in Table 1 and Supplementary Table 1.

**Table 1:**
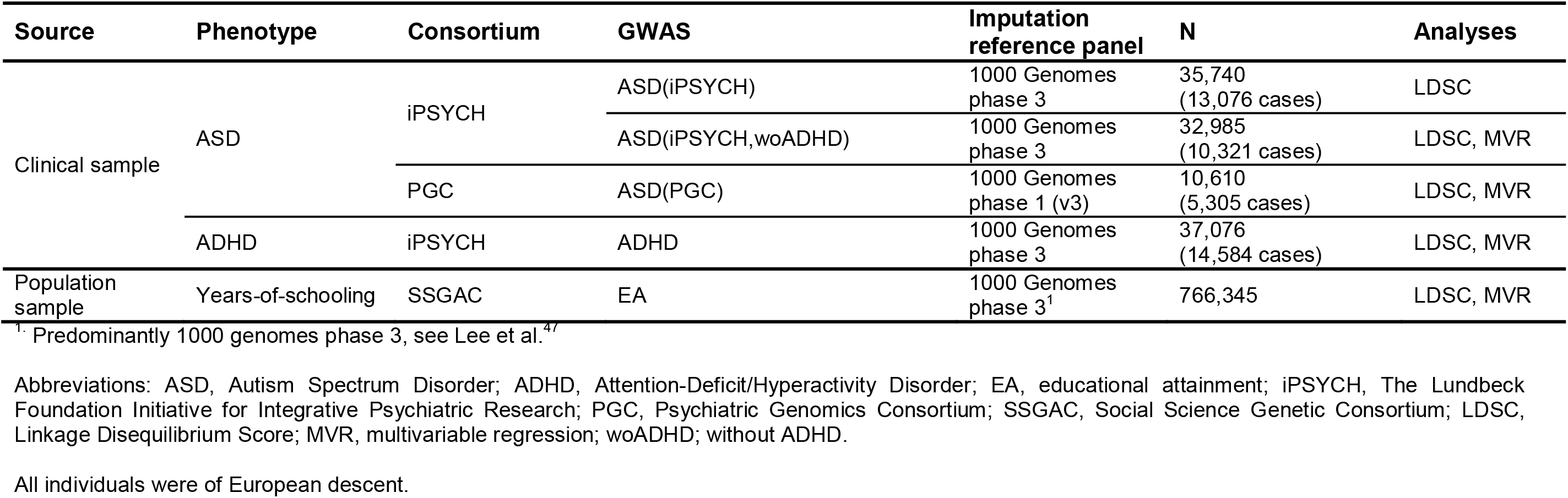
Sample description.

#### EA and general intelligence

GWAS summary statistics on years-of-schooling (discovery and replication sample excluding 23andMe) were obtained from the Social Science Genetic Association Consortium (SSGAC, https://www.thessgac.org/, Table 1)^47^. EA was coded according to the International Standard Classification of Education (1997) scale^50^ and analysed as a quantitative variable defined as an individual’s years-of-schooling. Participants were >30 years of age at the time of assessment and of European ancestry. The meta-analysis consisted primarily of population-based cohorts, but also included family-based and case-control samples. 55.2% of participants were female. For most cohorts, genome-wide data were imputed to a 1000 genomes project^51^ version 3 reference template, as previously described^47^.

GWAS summary statistics on general intelligence^27^ were retrieved from the Complex Trait Genetics (CTG) lab (https://ctg.cncr.nl/software/summary_statistics, Supplementary Table 1). Participating cohorts were primarily population-based. Each cohort assessed intelligence with different instruments that were re-defined to index a common latent factor of general intelligence (GI)^27^. Participants had a wide age range (from 5 to 98 years), 51.2% were female and all of them were of European descent. Genome-wide data were predominantly imputed to the Haplotype Reference Consortium (HRC) reference panel^52^, as previously described^27^.

#### ASD and ADHD

GWAS summary statistics for ASD and ADHD were accessed through the Danish Lundbeck Foundation Initiative for Integrative Psychiatric Research (iPSYCH, http://ipsych.au.dk/) using samples from the Danish Neonatal Screening Biobank hosted by Statens Serum Institute^21,28,53^ (ASD(iPSYCH), ADHD(iPSYCH), Table 1). iPSYCH adopts a case-control design (26.6% female ASD-cases^21^, 21.6% female ADHD-cases^28^) with shared controls (~49% female)^21,28^, all of European ancestry with age ranges spanning infancy to adulthood^21,29^. For MVR analyses, ASD samples were restricted to ASD-cases without (wo) an additional ADHD diagnosis (ASD(iPSYCH,woADHD), Table 1) to avoid overlap with ADHD(iPSYCH). However, ADHD-cases may have an additional ASD diagnosis. Information on ADHD cases without ASD was not available.

ASD cases and ADHD cases were diagnosed according to ICD-10^36^ and identified using the Danish Psychiatric Central Research Register^54^. Registry-based ASD diagnoses were validated previously^54,55^. Controls were randomly selected from the same nationwide birth cohort and did not have a diagnosis of ASD or ADHD or moderate-severe mental retardation (F71-F79)^28,53^. The median age at first diagnosis of ASD was 10 years. Genotyping was performed using the Illumina Infinium PsychArray BeadChip and genotypes were imputed to a 1000 Genomes template^51^ (Phase3, release 02-05-2013). Genotyping, quality control, imputation and genetic association analysis were carried out using the Ricopili pipeline with standard PGC settings^42^.

Independent ASD GWAS summary statistics were obtained from the Psychiatric Genomics Consortium (www.med.unc.edu/pgc/). They were based on a case-control/pseudo-control design and all individuals were ≥3 years of age and of European ancestry (ASD(PGC), Table 1). Information on the male-female ratio was not available^56^. A consensus ASD diagnosis was made using research standard diagnoses and expert clinical consensus diagnoses. The majority of ASD-cases (94.1%) also had a clinical diagnosis based on the Autism Diagnostic Interview-Revised^37^ and/or the Autism Diagnostic Observation Schedule^57^. Genome-wide data were imputed to a 1000 Genomes reference template^51^ (Phase1 v3).

For genetic correlation analyses, a combined ASD+ADHD GWAS statistic was derived by conducting a random-effect meta-analysis of ASD(PGC) and ADHD(iPSYCH) using GWAS summary statistics and GWAMA software^58^. Note that the sample size for ADHD(iPSYCH) is about three times larger than for ASD(PGC).

#### Other psychiatric disorders

To assess the specificity of MVR association profiles, we also investigated GWAS summary statistics for Major Depressive Disorder (MDD)^48^, Schizophrenia (SCZ)^49^ and Bipolar Disorder (BD)^49^. Cases were identified based on international consensus criteria. For MDD, cases were identified based on a lifetime diagnosis of MDD, established using DSM-III, DSM-IV, ICD-9 and/or ICD-10 criteria^48^. For SCZ, the majority of cases were diagnosed using DSM-III, DSM-III-R, DSM-IV, ICD-10, and SCID criteria^42,49^. BD cases were diagnosed according to DSM-III, DSM-IV-TR, DSM-IV, SCID, ICD-10 and/or RDC criteria^49,59^. For all three data sets, genotype imputation was performed using the IMPUTE2 / SHAPEIT pipeline against the 1000 Genomes Project (v3) template. Summary data were obtained from the PGC (www.med.unc.edu/pgc/, Supplementary Table 1), all based on participants of European ancestry.

### SNP-heritability and genetic correlations

SNP-heritability (SNP-h^2^), as the proportion of phenotypic or liability variance tagged by SNPs on genotyping arrays, was estimated for EA, general intelligence and psychiatric disorders using Linkage Disequilibrium Score (LDSC) regression^60^ (Supplementary Table 2). To estimate LDSC-h^2^, genome-wide χ^2^-statistics are regressed on the amount of genetic variation captured by each Single Nucleotide Polymorphism (SNP)^60^, while the intercept of this regression minus one is an estimator of the mean contribution of confounding bias to the inflation in the mean χ^2^-statistic^60^. SNP-h^2^ was calculated on the liability scale for psychiatric disorder samples, assuming a population prevalence of 0.012 for ASD^21^, 0.05 for ADHD^61^, 0.162 for Major Depressive Disorder (MDD)^62^, 0.007 for Schizophrenia (SCZ)^63^ and 0.006 for Bipolar Disorder (BD)^64^.

In extension, unconstrained LDSC correlation^65^ analysis was applied to estimate bivariate genetic correlations (r_g_) (Supplementary Table 3-5), as a regression of the product of test statistics on LD score that captures the extent of shared genetic influences between phenotypes assessed in distinct samples. All analyses were performed with LDSC software^65,66^ and based on the set of well-imputed HapMap3 SNPs and a European reference panel of LD scores^65^.

### Multivariable regression analysis

We conducted a set of interlinked MVR analyses to dissect polygenic ASD-EA and ADHD-EA associations into either ASD-specific or ADHD-specific associations as well as genetic influences that are shared across both disorders and EA (overview in Supplementary Figure 1). Here, MVR (sometimes also referred to as multiple linear regression) contains a single outcome (dependent variable) and multiple predictors (independent variables). Specifically, we translated a causal modelling approach using GWAS summary statistics^32^ into a polygenic context without making causal inferences. In principle, this involves a weighted multivariable regression approach, where we regressed SNP estimates for EA (Z_EA,_ dependent variable) jointly on SNP estimates for ASD (X_ASD_, independent variable) and SNP estimates for ADHD (Y_ADHD_, independent variable)(Formula 1-4). This methodology^32^ can disentangle polygenetic trait interrelationships using single SNP information, while controlling for bias^67^ and assesses here, due to the polygenic context, genetic associations only.

#### Genetic variant selection

For all discovery ASD- and ADHD-MVR analyses (Supplementary Figure 1a), 11 ASD-related and 11 ADHD-related variant sets were selected from ASD(iPSYCH,woADHD) and ADHD(iPSYCH) GWAS statistics respectively, using multiple *P*-value thresholds (*P_thr_*, 5×10^−8^; 5×10^−7^; 5×10^−6^; 5×10^−5^; 0.0005; 0.0015; 0.005; 0.05; 0.1; 0.3; 0.5), similar to a polygenic scoring approach^31^. However, for presented MVR analyses (including sensitivity and follow-up analyses) we predominantly focused on two *P*-value thresholds only: (i) *P*_thr_<0.0015, consistent with guidelines for validating genetic instrument strength (F-statistic<10)^68^ and conservative selection thresholds recommended for related polygenic scoring approaches^31^, and (ii) *P*_thr_<0.05, a less stringent threshold that has been previously applied for the polygenic analysis of complex psychiatric disorders^42^, with the aim of increasing the statistical power and precision of MVR estimates. All variants were restricted to common (minor allele frequency>0.01), independent (linkage disequilibrium-r^2^<0.25 within ±500 kb) and well-imputed (Imputation quality(INFO) >0.7) SNPs.

#### Estimation of ASD-specific, ADHD-specific and cross-disorder genetic associations with EA

After creating variant sets for both ASD and ADHD, SNP estimates for these variants were extracted from ASD(iPSYCH, woADHD), ADHD(iPSYCH) and EA(SSGAC) GWAS statistics. Next, per ASD variant set, an ASD-MVR was fitted as follows (Formula 1-2):

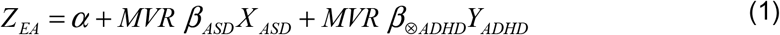

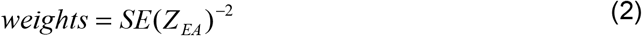

with SNP estimates Z_EA_ (independent variable), X_ASD_ (dependent variable) and Y_ADHD_ (dependent variable), regression intercept α, ASD-specific MVR effect β_ASD_, and cross-disorder MVR effect β_⊗ADHD_. Thus, in ASD-MVR models (Supplementary Figure 1a), ASD-specific associations with EA (ASD-MVR β_ASD_) were assessed using ASD SNP estimates and cross-disorder associations (MVR β_⊗ADHD_) with EA using ADHD GWAS estimates.

Similarly, per ADHD variant set, an ADHD-MVR was fitted as (Formula 3-4):

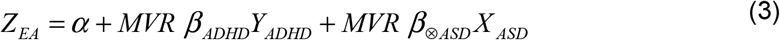

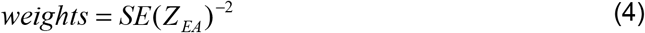

with SNP estimates Z_EA_ (independent variable), X_ASD_ (dependent variable) and Y_ADHD_ (dependent variable), regression intercept α, ADHD-specific MVR effect β_ADHD_, and cross-disorder MVR effect MVR β_⊗ASD_. Here, in ADHD-MVR (Supplementary Figure 1a), ADHD-specific associations with EA (ADHD-MVR β_ADHD_) were assessed using ADHD SNP estimates and cross-disorder associations with EA (MVR β_⊗ASD_) using ASD SNP estimates.

Cross-disorder effects were thus estimated twice: (i) fitting ADHD SNP estimates for ASD variant sets and (ii) fitting ASD SNP estimates for ADHD variant sets.

Reported MVR effects present changes in years-of-schooling per increase in log-odds ASD or ADHD liability, respectively, pooled across the selected variants. The overall model fit of MVRs was compared to univariable regression models (see below) using likelihood-ratio tests, as implemented in the R:stats library (Rv3.5.1). To assess collinearity between predictors, we calculated the variance inflation factor (VIF, R:car library (Rv3.5.1)).

All MVR models included an intercept (α) to allow for the presence of alternative pathways between ASD- and ADHD-related genetic variants and EA, other than captured by ASD or ADHD SNP estimates (unconstrained MVR models). An intercept consistent with zero (i.e. within the 95% confidence interval) suggests that there is no evidence for additional pleiotropic effects. As MVR estimates are thus sensitive to allelic alignment, MVR models were fitted twice: (i) a discovery MVR with SNP effects aligned to ASD risk in ASD-MVRs, and ADHD risk in ADHD-MVRs, respectively (Supplementary Figure 1a, across 11 *P*-value thresholds described above (5×10^−8^≤*P*_thr_<0.5); (ii) a subset of variants from (i) with concordant association effects for both ASD and ADHD risk (Supplementary Figure 1b, *P*-value thresholds: *P*_thr_<0.0015; *P*_thr_<0.5).

To replicate MVR findings from the discovery analysis, both ASD-MVR and ADHD-MVR findings were followed-up using ASD(PGC) instead of ASD(iPSYCH,woADHD) SNP estimates and the set of overlapping SNPs between ASD(iPSYCH,woADHD) and ASD(PGC) (Supplementary Figure 1c, *P*-value thresholds: *P*_thr_<0.0015; *P*_thr_<0.05). In addition, ASD-MVR and ADHD-MVR models were fitted using summary statistics for general intelligence, instead of EA (Supplementary Figure 1d, *P*-value thresholds: *P*_thr_<0.0015; *P*_thr_<0.05).

To identify loci underlying the observed MVR cross-disorder associations, we restricted ASD-MVR and ADHD-MVR variant sets at *P_th_*_r_<0.0015 (ASD: N_SNPs_≤1,973, ADHD: N_SNPs_≤2,717) to SNPs that were associated with both disorders at various *P*-value thresholds, and then re-analysed them (Supplementary Figure 1e). For this, we assessed the proportion of overlapping independent SNPs associated with both ASD and ADHD risk using PLINK (500 kb and Linkage Disequilibrium-r^2^≥0.6). We started with two variant sets of interest: (1) ASD-related variants at *P*_thr_<0.0015 (N_SNPs_=1,973) and (2) ADHD-related variants at *P*_thr_<0.0015 (N_SNPs_ =2,717). For each variant set, we identified the percentage of SNPs that were also associated with the other disorder across a range of *P*-value thresholds (5×10^−8^; 5×10^−7^; 5×10^−6^; 5×10^−5^; 0.0005; 0.0015; 0.005; 0.05; 0.1; 0.3; 0.5). In total, this resulted in 11 subsets for ASD-related variants, and 11 subsets for ADHD-related variants. Next, we performed MVRs using these SNP subsets only. For variant subsets with the largest MVR effects, we identified the corresponding genes based on positional mapping using PLINK software (0 kb gene window), similar to the default options applied by current gene-enrichment software^69^. Gene positions were retrieved from UCSC RefSeq gene range lists (genome build 37).

To assess the specificity of observed ASD/ADHD cross-disorder associations with EA, we carried out sensitivity MVR analyses that were similar to the discovery analyses described above. We used the same ASD and ADHD variants sets for ASD-MVR and ADHD-MVR, respectively, but replaced the SNP estimates for the cross-disorder (i.e. ADHD in ASD-MVR and ASD in ADHD-MVR)with SNP estimates from MDD, SCZ or BD GWAS (Supplementary Table 1, Supplementary Figure 1f, *P*-value thresholds: *P*_thr_<0.0015; *P*_thr_<0.05).

We applied the following conservative Bonferroni-corrected multiple testing thresholds for the MVR analyses described above: (i) discovery analyses with two MVR models across 11 *P-*value thresholds (22 tests, *P*_*Adjusted*_=0.0023, Supplementary Figure 1a) with concordant SNP set analyses being nested within these discovery analyses (Supplementary Figure 1b); (ii) follow-up analyses with independent ASD(PGC) estimates with two MVR models across two *P*-value thresholds (4 tests, *P*_*Adjusted*_=0.0125, Supplementary Figure 1c); (iii) follow-up analyses with independent general intelligence (CTG) estimates with two MVR models across two *P*-value thresholds (4 tests, *P*_*Adjusted*_=0.0125, Supplementary Figure 1d); (iv) screening of MVR effect sizes with variant sets, associated with both ASD and ADHD risk, applying joint ASD and ADHD variant selection criteria: For each MVR model, variant sets selected at *P*_*th*r_<0.0015 were successively restricted to SNPs that are associated with both disorders, using 11 *P*-value thresholds (2 × MVR models × 11 tests + 22 additional tests correcting for all discovery analyses, 44 total tests, *P*_*Adjusted*_=0.0011, Supplementary Figure 1e) and (v) follow-up analyses with independent MDD(PGC), SCZ(PGC) and BD(PGC) estimates with two MVR models across two *P*-value thresholds (12 tests, *P*_*Adjusted*_=0.0042, Supplementary Figure 1f).

### Univariable regression models

For comparison with MVR, univariable weighted regression models were conducted based on ASD and ADHD variant sets (*P*_thr_<0.0015 and *P*_thr_<0.05) selected from ASD(iPSYCH, woADHD) and ADHD(iPSYCH) GWAS statistics, respectively. Corresponding SNP estimates for ASD, ADHD and EA were subsequently extracted from ASD(iPSYCH, woADHD), ADHD(iPSYCH) and EA(SSGAC) GWAS statistics, as described for MVR above.

Using univariable weighted regression models (R:stats library, Rv3.5.1) and ASD variants, (1) SNP estimates for ADHD were regressed on SNP estimates for ASD, (2) SNP estimates for EA were regressed on SNP estimates for ASD, and (3) SNP estimates for EA were regressed on SNP estimates for ADHD. Similar models were fitted using ADHD variants. All univariable regressions included an intercept. Models were fitted twice: (i) with SNP estimates aligned according to the risk-increasing allele for the disorder used for variant selection and (ii) a subset of variants from (i) with concordant association effect for ASD and ADHD risk. The univariable model fit was compared with MVRs using a likelihood-ratio test as implemented in the R:stats library (Rv3.5.1).

### Structural equation modelling

To summarise genetic interrelationships between EA, ASD and ADHD with a multi-factor model, we translated known SNP-h^2^ and r_g_ estimates (Supplementary Table 3-4) into hypothetical factor loadings consistent with structural equations for a saturated model. Specifically, we propose a multi-factorial structural equation model consisting of three continuous phenotypes (EA, ASD liability and ADHD liability Z-scores), three independent latent genetic factors, and three independent latent residual influences. We assume that genetic factors give rise to genetic variances and covariances between EA, ASD and ADHD liability, while residual covariances are assumed to be absent. Phenotypic variances and covariances were described according to a Cholesky decomposition^70^ (i.e. a saturated model), assuming an infinitely large population and a fully identified model (Figure 3, Supplementary Figure 4). A Cholesky model involves the decomposition of both the genetic variances and residual variances into as many latent factors as there are observed variables. The expected phenotypic covariance matrix Σ for Z-standardised traits based on the factor model is

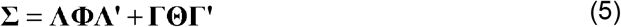

where Λ is a lower triangular matrix of genetic factor loadings, Φ is a diagonal matrix of latent genetic factor variances (standardised to unit variance) such that Φ is an identity matrix ***I***. The residual variance can be decomposed into latent residual factors, where Γ is a lower triangular matrix of residual factor loadings and Θ is a diagonal matrix of latent residual factor variances (standardised to unit variance) such that Θ is an identity matrix ***I***. For example, for a trivariate model consisting of measures P_1_, P_2_ and P_3_, assuming three genetic factors (A_1_, A_2_ and A_3_) and three residual factors (E_1_, E_2_ and E_3_), the expected phenotypic covariance matrix can be expressed as follows:

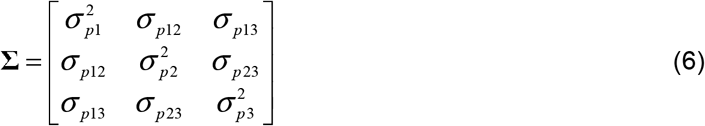

with the relevant matrices

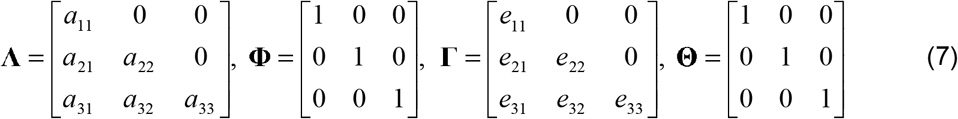

where 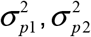 and 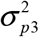 represent the phenotypic variances and *σ*_*p*12_, *σ*_*p*13_ and *σ*_*p*23_ phenotypic covariances. We annotate the genetic factor loadings *a* (factor loadings) such that the first number indicates the direction of the effect (the variable to which the arrow points) and the second the origin of the effect.

The trivariate AE Cholesky decomposition of three standardised measures, as described above, can be visualised by means of a path diagram (Supplementary Figure 4) and the expected phenotypic variances and covariances can be expressed as follows:

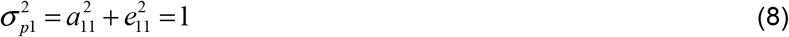

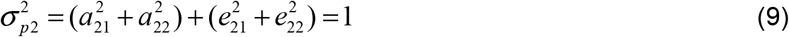

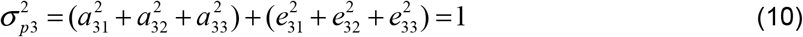

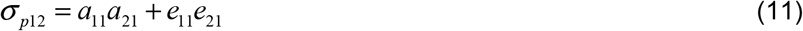

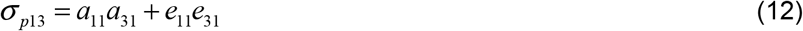

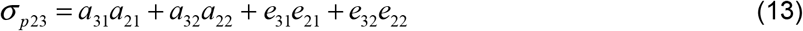

The variance of the latent genetic and residual factors has been standardised to unit variance and is not shown.

Estimated genetic variances and covariances can be used to derive genetic correlation estimates between two phenotypes measuring the extent to which two phenotypes 1 and 2 share genetic factors (ranging from −1 to 1):

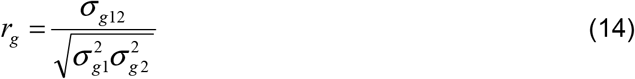

where *σ*_*g*12_ is the genetic covariance between phenotypes 1 and 2 and 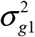 and 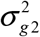 the genetic variances.

We derived (but did not fit) hypothetical factor loadings, based on LDSC SNP-h^2^ estimates for EA and, on the liability scale, ASD and ADHD (Supplementary Table 2), as well as unconstrained LDSC genetic correlations (Supplementary Table 3-4) using EA, ASD(iPSYCH), ASD(PGC) and ADHD(iPSYCH) GWAS statistics (Table 1). We support the plausibility of such a model using simulations (Supplementary Table 18).

### Multi-factor model data simulation

To evaluate the accuracy of the proposed multifactorial model, we carried out data simulations (Supplementary Table 18). Assuming multivariate normality and unrelated individuals, we simulated three continuous interrelated measures P_1_, P_2_ and P_3_ corresponding to EA and liability for ADHD and ASD respectively, assuming an underlying Cholesky model. This includes three genetic factors with their variances and covariances and three residual factors with their variances and their covariances. The genetic interrelationships between these three traits were informed by unconstrained LDSC genetic correlations between EA, ADHD and ASD (Supplementary Table 3-4) using EA, ADHD(iPSYCH) and ASD(PGC) summary statistics, using structural equations described above (8 to 14). Residual interrelationships were assumed to be absent as the cohorts are independent of each other. However, simulated SNP-h^2^ estimates were increased, and sample size restricted to 6,000 individuals per trait with 20,000 SNPs per genetic factor, to ease the computational burden (72h, using 16 cores). Multivariate variances and covariances within the simulated data were modelled using genetic-relationship structural equation modelling (GSEM, R gsem library, v0.1.2)^71^. This method involves a multivariate analysis of genetic variance by combining whole-genome genotyping information in unrelated individuals with structural equation modelling techniques using a full information maximum likelihood approach. Simulated parameters and estimated parameters are shown in Supplementary Table 18.

## Data availability

Genome-wide association summary statistics on ASD(PGC), EA(SSGAC), GI(CTG), MDD(PGC), SCZ(PGC) and BD(PGC) are publically available. Download links are provided in the methods section. Restrictions apply to the availability of summary statistics from the iPSYCH sample. For access to these data, researchers should contact the lead principal investigator A.D.B.

## Acknowledgements

This research was facilitated by the Social Science Genetic Association Consortium (SSGAC), Complex Trait Genetics (CTG) lab, Psychiatric Genomics Consortium (PGC) and iPSYCH-Broad-PGC ASD Consortium, by providing access to genome-wide summary statistics.

This publication is the work of the authors and EV and BSTP will serve as guarantors for the contents of this paper. EV and BSTP are supported by the Max Planck Society. BSTP is supported by the Simons Foundation (Award ID: 514787). The iPSYCH project is funded by the Lundbeck Foundation (grant no R102-A9118 and R155-2014-1724) and the universities and university hospitals of Aarhus and Copenhagen. ADB is also supported by the EU’s Horizon 2020 programme (grant no 667302, CoCA). Data handling and analysis was supported by NIMH (1U01MH109514-01 to Michael O’Donovan and Anders D Børglum). High-performance computer capacity for handling and statistical analysis of iPSYCH data on the GenomeDK HPC facility was provided by the Centre for Integrative Sequencing, iSEQ, Aarhus University, Denmark (grant to Anders D Børglum) and Center for Genomics and Personalized Medicine, Aarhus, Denmark. We thank Simon E. Fisher for helpful discussions of the manuscript.

## Author contributions

B.S.P. and A.D.B. designed and supervised the research. E.V., J.G. and D.D. analysed the genetic data. C.Y.S. and S.B provided methodological support. E.V., J.G. and B.S.P. wrote the manuscript. All authors read and commented on the manuscript.

## References

1. Thapar A, Cooper M. Attention deficit hyperactivity disorder. The Lancet. 2016;387(10024):1240–1250. doi:10.1016/S0140-6736(15)00238-X

2. Lord C, Elsabbagh M, Baird G, Veenstra-Vanderweele J. Autism spectrum disorder. The Lancet. 2018;392(10146):508–520. doi:10.1016/S0140-6736(18)31129-2

3. Antshel KM, Zhang-James Y, Wagner KE, Ledesma A, Faraone SV. An update on the comorbidity of ADHD and ASD: a focus on clinical management. Expert Rev Neurother. 2016;16(3):279–293. doi:10.1586/14737175.2016.1146591

4. Stergiakouli E, Hamshere M, Holmans P, et al. Investigating the contribution of common genetic variants to the risk and pathogenesis of ADHD. Am J Psychiatry. 2012;169(2):186–194. doi:10.1176/appi.ajp.2011.11040551

5. Gaugler T, Klei L, Sanders SJ, et al. Most genetic risk for autism resides with common variation. Nat Genet. 2014;46(8):881–885. doi:10.1038/ng.3039

6. Krumm N, Turner TN, Baker C, et al. Excess of rare, inherited truncating mutations in autism. Nat Genet. 2015;47(6):582–588. doi:10.1038/ng.3303

7. Williams NM, Zaharieva I, Martin A, et al. Rare chromosomal deletions and duplications in attention-deficit hyperactivity disorder: a genome-wide analysis. Lancet Lond Engl. 2010;376(9750):1401–1408. doi:10.1016/S0140-6736(10)61109-9

8. Rommelse NNJ, Franke B, Geurts HM, Hartman CA, Buitelaar JK. Shared heritability of attention-deficit/hyperactivity disorder and autism spectrum disorder. Eur Child Adolesc Psychiatry. 2010;19(3):281–295. doi:10.1007/s00787-010-0092-x

9. Leitner Y. The co-occurrence of autism and attention deficit hyperactivity disorder in children - what do we know? Front Hum Neurosci. 2014;8:268. doi:10.3389/fnhum.2014.00268

10. Ronald A, Simonoff E, Kuntsi J, Asherson P, Plomin R. Evidence for overlapping genetic influences on autistic and ADHD behaviours in a community twin sample. J Child Psychol Psychiatry. 2008;49(5):535–542. doi:10.1111/j.1469-7610.2007.01857.x

11. Stergiakouli E, Davey Smith G, Martin J, et al. Shared genetic influences between dimensional ASD and ADHD symptoms during child and adolescent development. Mol Autism. 2017;8:18. doi:10.1186/s13229-017-0131-2

12. Ronald A, Larsson H, Anckarsäter H, Lichtenstein P. Symptoms of autism and ADHD: A Swedish twin study examining their overlap. J Abnorm Psychol. 2014;123(2):440–451. doi:10.1037/a0036088

13. Polderman TJC, Hoekstra RA, Posthuma D, Larsson H. The co-occurrence of autistic and ADHD dimensions in adults: an etiological study in 17◻770 twins. Transl Psychiatry. 2014;4(9):e435. doi:10.1038/tp.2014.84

14. Taylor MJ, Charman T, Ronald A. Where are the strongest associations between autistic traits and traits of ADHD? evidence from a community-based twin study. Eur Child Adolesc Psychiatry. 2015;24(9):1129–1138. doi:10.1007/s00787-014-0666-0

15. Taylor MJ, Charman T, Robinson EB, et al. Developmental associations between traits of autism spectrum disorder and attention deficit hyperactivity disorder: a genetically informative, longitudinal twin study. Psychol Med. 2013;43(8):1735–1746. doi:10.1017/S003329171200253X

16. Martin J, Hamshere ML, Stergiakouli E, O’Donovan MC, Thapar A. Genetic Risk for Attention-Deficit/Hyperactivity Disorder Contributes to Neurodevelopmental Traits in the General Population. Biol Psychiatry. 2014;76(8):664–671. doi:10.1016/j.biopsych.2014.02.013

17. Lichtenstein P, Carlström E, Råstam M, Gillberg C, Anckarsäter H. The Genetics of Autism Spectrum Disorders and Related Neuropsychiatric Disorders in Childhood. Am J Psychiatry. 2010;167(11):1357–1363. doi:10.1176/appi.ajp.2010.10020223

18. Ghirardi L, Brikell I, Kuja-Halkola R, et al. The familial co-aggregation of ASD and ADHD: a register-based cohort study. Mol Psychiatry. 2018;23(2):257–262. doi:10.1038/mp.2017.17

19. Martin J, Cooper M, Hamshere ML, et al. Biological overlap of attention-deficit/hyperactivity disorder and autism spectrum disorder: evidence from copy number variants. J Am Acad Child Adolesc Psychiatry. 2014;53(7):761–770.e26. doi:10.1016/j.jaac.2014.03.004

20. Martin J, Taylor MJ, Lichtenstein P. Assessing the evidence for shared genetic risks across psychiatric disorders and traits. Psychol Med. 2018;48(11):1759–1774. doi:10.1017/S0033291717003440

21. Grove J, Ripke S, Als TD, et al. Identification of common genetic risk variants for autism spectrum disorder. Nat Genet. 2019;51(3):431. doi:10.1038/s41588-019-0344-8

22. Cross-Disorder Group of the Psychiatric Genomics Consortium. Genetic relationship between five psychiatric disorders estimated from genome-wide SNPs. Nat Genet. 2013;45(9):984–994. doi:10.1038/ng.2711

23. Bulik-Sullivan B, Finucane HK, Anttila V, et al. An atlas of genetic correlations across human diseases and traits. Nat Genet. 2015;47(11):1236–1241. doi:10.1038/ng.3406

24. Okbay A, Beauchamp JP, Fontana MA, et al. Genome-wide association study identifies 74 loci associated with educational attainment. Nature. 2016;533(7604):539–542. doi:10.1038/nature17671

25. Martin J, Hamshere ML, Stergiakouli E, O’Donovan MC, Thapar A. Neurocognitive abilities in the general population and composite genetic risk scores for attention-deficit hyperactivity disorder. J Child Psychol Psychiatry. 2015;56(6):648–656. doi:10.1111/jcpp.12336

26. Stergiakouli E, Martin J, Hamshere ML, et al. Association between polygenic risk scores for attention-deficit hyperactivity disorder and educational and cognitive outcomes in the general population. Int J Epidemiol. 2017;46(2):421–428. doi:10.1093/ije/dyw216

27. Savage JE, Jansen PR, Stringer S, et al. Genome-wide association meta-analysis in 269,867 individuals identifies new genetic and functional links to intelligence. Nat Genet. 2018;50(7):912–919. doi:10.1038/s41588-018-0152-6

28. Demontis D, Walters RK, Martin J, et al. Discovery of the first genome-wide significant risk loci for attention deficit/hyperactivity disorder. Nat Genet. November 2018:1. doi:10.1038/s41588-018-0269-7

29. Clarke T-K, Lupton MK, Fernandez-Pujals AM, et al. Common polygenic risk for autism spectrum disorder (ASD) is associated with cognitive ability in the general population. Mol Psychiatry. 2016;21(3):419–425. doi:10.1038/mp.2015.12

30. Selzam S, Coleman JRI, Caspi A, Moffitt TE, Plomin R. A polygenic p factor for major psychiatric disorders. Transl Psychiatry. 2018;8(1):205. doi:10.1038/s41398-018-0217-4

31. Wray NR, Lee SH, Mehta D, Vinkhuyzen AAE, Dudbridge F, Middeldorp CM. Research Review: Polygenic methods and their application to psychiatric traits. J Child Psychol Psychiatry. 2014;55(10):1068–1087. doi:10.1111/jcpp.12295

32. Burgess S, Thompson DJ, Rees JMB, Day FR, Perry JR, Ong KK. Dissecting Causal Pathways Using Mendelian Randomization with Summarized Genetic Data: Application to Age at Menarche and Risk of Breast Cancer. Genetics. January 2017:genetics.300191.2017. doi:10.1534/genetics.117.300191

33. Watanabe K, Stringer S, Frei O, et al. A global overview of pleiotropy and genetic architecture in complex traits. Nat Genet. August 2019:1–10. doi:10.1038/s41588-019-0481-0

34. Satterstrom FK, Walters RK, Singh T, et al. ASD and ADHD have a similar burden of rare protein-truncating variants. bioRxiv. March 2018:277707. doi:10.1101/277707

35. Association AP. Diagnostic and Statistical Manual of Mental Disorders, 5th Edition: DSM-5. 5 edition. Washington, D.C: American Psychiatric Publishing; 2013.

36. International Statistical Classification of Diseases and Related Health Problems. Vol 10th revision. Malta: World Health Organization; 2010.

37. Lord C, Rutter M, Le Couteur A. Autism Diagnostic Interview-Revised: a revised version of a diagnostic interview for caregivers of individuals with possible pervasive developmental disorders. J Autism Dev Disord. 1994;24(5):659–685.

38. Boyle EA, Li YI, Pritchard JK. An Expanded View of Complex Traits: From Polygenic to Omnigenic. Cell. 2017;169(7):1177–1186. doi:10.1016/j.cell.2017.05.038

39. Liu X, Li YI, Pritchard JK. Trans Effects on Gene Expression Can Drive Omnigenic Inheritance. Cell. 2019;177(4):1022–1034.e6. doi:10.1016/j.cell.2019.04.014

40. Morris KV, Mattick JS. The rise of regulatory RNA. Nat Rev Genet. 2014;15(6):423–437. doi:10.1038/nrg3722

41. Gonda X, Hullam G, Antal P, et al. Significance of risk polymorphisms for depression depends on stress exposure. Sci Rep. 2018;8(1):3946. doi:10.1038/s41598-018-22221-z

42. Schizophrenia Working Group of the Psychiatric Genomics Consortium, Ripke S, Neale BM, et al. Biological insights from 108 schizophrenia-associated genetic loci. Nature. 2014;511(7510):421–427. doi:10.1038/nature13595

43. Mayes SD, Calhoun SL, Mayes RD, Molitoris S. Autism and ADHD: Overlapping and discriminating symptoms. Res Autism Spectr Disord. 2012;6(1):277–285. doi:10.1016/j.rasd.2011.05.009

44. Burgess S, Davies NM, Thompson SG. Bias due to participant overlap in two◻sample Mendelian randomization. Genet Epidemiol. 2016;40(7):597–608. doi:10.1002/gepi.21998

45. Wray NR, Yang J, Hayes BJ, Price AL, Goddard ME, Visscher PM. Pitfalls of predicting complex traits from SNPs. Nat Rev Genet. 2013;14(7):507–515. doi:10.1038/nrg3457

46. Greven CU, Harlaar N, Dale PS, Plomin R. Genetic Overlap between ADHD Symptoms and Reading is largely Driven by Inattentiveness rather than Hyperactivity-Impulsivity. J Can Acad Child Adolesc Psychiatry. 2011;20(1):6–14.

47. Lee JJ, Wedow R, Okbay A, et al. Gene discovery and polygenic prediction from a genome-wide association study of educational attainment in 1.1 million individuals. Nat Genet. 2018;50(8):1112–1121. doi:10.1038/s41588-018-0147-3

48. Wray NR, Ripke S, Mattheisen M, et al. Genome-wide association analyses identify 44 risk variants and refine the genetic architecture of major depression. Nat Genet. 2018;50(5):668. doi:10.1038/s41588-018-0090-3

49. Ruderfer DM, Ripke S, McQuillin A, et al. Genomic Dissection of Bipolar Disorder and Schizophrenia, Including 28 Subphenotypes. Cell. 2018;173(7):1705–1715.e16. doi:10.1016/j.cell.2018.05.046

50. Rietveld CA, Medland SE, Derringer J, et al. GWAS of 126,559 individuals identifies genetic variants associated with educational attainment. Science. 2013;340(6139):1467–1471. doi:10.1126/science.1235488

51. The 1000 Genomes Project Consortium. A global reference for human genetic variation. Nature. 2015;526(7571):68–74. doi:10.1038/nature15393

52. McCarthy S, Das S, Kretzschmar W, Durbin R, Abecasis G, Marchini J. A reference panel of 64,976 haplotypes for genotype imputation. bioRxiv. April 2016:035170. doi:10.1101/035170

53. Pedersen CB, Bybjerg-Grauholm J, Pedersen MG, et al. The iPSYCH2012 case-cohort sample: new directions for unravelling genetic and environmental architectures of severe mental disorders. Mol Psychiatry. September 2017. doi:10.1038/mp.2017.196

54. Mors O, Perto GP, Mortensen PB. The Danish Psychiatric Central Research Register. Scand J Public Health. 2011;39(7 Suppl):54–57. doi:10.1177/1403494810395825

55. Lauritsen MB, Jørgensen M, Madsen KM, et al. Validity of childhood autism in the Danish Psychiatric Central Register: findings from a cohort sample born 1990-1999. J Autism Dev Disord. 2010;40(2):139–148. doi:10.1007/s10803-009-0818-0

56. Cross-Disorder Group of the Psychiatric Genomics Consortium. Identification of risk loci with shared effects on five major psychiatric disorders: a genome-wide analysis. Lancet Lond Engl. 2013;381(9875):1371–1379. doi:10.1016/S0140-6736(12)62129-1

57. Lord C, Risi S, Lambrecht L, et al. The autism diagnostic observation schedule-generic: a standard measure of social and communication deficits associated with the spectrum of autism. J Autism Dev Disord. 2000;30(3):205–223.

58. Mägi R, Morris AP. GWAMA: software for genome-wide association meta-analysis. BMC Bioinformatics. 2010;11:288. doi:10.1186/1471-2105-11-288

59. Stahl EA, Breen G, Forstner AJ, et al. Genome-wide association study identifies 30 loci associated with bipolar disorder. Nat Genet. 2019;51(5):793–803. doi:10.1038/s41588-019-0397-8

60. Bulik-Sullivan BK, Loh PR, Finucane HK, et al. LD Score regression distinguishes confounding from polygenicity in genome-wide association studies. Nat Genet. 2015;47(3):291–295. doi:10.1038/ng.3211

61. Polanczyk G, de Lima MS, Horta BL, Biederman J, Rohde LA. The Worldwide Prevalence of ADHD: A Systematic Review and Metaregression Analysis. Am J Psychiatry. 2007;164(6):942–948. doi:10.1176/ajp.2007.164.6.942

62. Kupfer DJ, Frank E, Phillips ML. Major depressive disorder: new clinical, neurobiological, and treatment perspectives. The Lancet. 2012;379(9820):1045–1055. doi:10.1016/S0140-6736(11)60602-8

63. Owen MJ, Sawa A, Mortensen PB. Schizophrenia. The Lancet. 2016;388(10039):86–97. doi:10.1016/S0140-6736(15)01121-6

64. Merikangas KR, Jin R, He J-P, et al. Prevalence and correlates of bipolar spectrum disorder in the world mental health survey initiative. Arch Gen Psychiatry. 2011;68(3):241–251. doi:10.1001/archgenpsychiatry.2011.12

65. Bulik-Sullivan B, Finucane HK, Anttila V, et al. An atlas of genetic correlations across human diseases and traits. Nat Genet. 2015;47:1236–1241. doi:10.1038/ng.3406

66. Bulik-Sullivan BK, Loh P-R, Finucane HK, et al. LD Score regression distinguishes confounding from polygenicity in genome-wide association studies. Nat Genet. 2015;47(3):291–295. doi:10.1038/ng.3211

67. Aschard H, Vilhjálmsson BJ, Joshi AD, Price AL, Kraft P. Adjusting for heritable covariates can bias effect estimates in genome-wide association studies. Am J Hum Genet. 2015;96(2):329–339. doi:10.1016/j.ajhg.2014.12.021

68. Pierce BL, Ahsan H, Vanderweele TJ. Power and instrument strength requirements for Mendelian randomization studies using multiple genetic variants. Int J Epidemiol. 2011;40(3):740–752. doi:10.1093/ije/dyq151

69. Leeuw CA de, Mooij JM, Heskes T, Posthuma D. MAGMA: Generalized Gene-Set Analysis of GWAS Data. PLOS Comput Biol. 2015;11(4):e1004219. doi:10.1371/journal.pcbi.1004219

70. Neale M, Boker S, Xie G, Maes HHM. Mx: Statistical Modeling. 7th ed. Richmond: Department of Psychiatry; 2006.

71. St Pourcain B, Eaves LJ, Ring SM, et al. Developmental Changes Within the Genetic Architecture of Social Communication Behavior: A Multivariate Study of Genetic Variance in Unrelated Individuals. Biol Psychiatry. 2018;83(7):598–606. doi:10.1016/j.biopsych.2017.09.020

