## Supplementary Information for "Shared risk alleles with discordant polygenic effects: Disentangling the genetic overlap between ASD and ADHD"

### Supplementary Methods

#### Web resources

SSGAC: https://www.thessgac.org/

CTG lab: https://ctg.cncr.nl/software/summary_statistics

PGC: https://www.med.unc.edu/pgc/

iPSYCH: https://ipsych.au.dk/downloads/

PLINK: https://www.cog-genomics.org/plink2

LDSC: https://github.com/bulik/ldsc

R: https://www.r-project.org/

### Supplementary Tables

Supplementary Table 1: Sample description for MDD, SCZ, BD and general intelligence

| Source | Phenotype | Consortium | Imputation reference panel | N | Analyses |
|---|---|---|---|---|---|
| Clinical sample | MDD | PGC | 1000 Genomes multi-ancestry | 173,005 (cases=59,851) | LDSC, MVR |
| | SCZ | PGC | 1000 Genomes phase 3 | 65,967 (cases=33,426) | LDSC, MVR |
| | BD | PGC | 1000 Genomes phase 3 | 41,653 (cases=20,129) | LDSC, MVR |
| Population sample | GI | CTG lab | HRC[1] | 279,930 | LDSC, MVR |

[1] Predominantly HRC, see Savage et al.[1]

Abbreviations: BD, Bipolar Disorder; CTG, Complex Trait Genetics lab; GI, general intelligence; HRC, Haplotype Reference Consortium; LDSC, Linkage Disequilibrium Score regression and correlation; MDD, Major Depressive Disorder; MVR, multivariable regression; PGC, Psychiatric Genomics Consortium; SCZ, Schizophrenia

All individuals were of European descent.

Supplementary Table 2: SNP-heritability estimates

| Phenotype | Sample | SNP-$h^2$ (SE) | $\lambda_{GC}$ | Intercept (SE) |
|---|---|---|---|---|
| ASD | ASD(iPSYCH) | 0.12(0.01) | 1.15 | 1.01(0.01) |
| | ASD(iPSYCH, woADHD) | 0.13(0.01) | 1.13 | 1.01(0.01) |
| | ASD(PGC) | 0.26(0.03) | 1.05 | 0.97(0.01) |
| ADHD | ADHD(iPSYCH) | 0.26(0.02) | 1.23 | 1.03(0.01) |
| MDD | MDD(PGC) | 0.095(0.01) | 1.24 | 0.99(0.01) |
| SCZ | SCZ(PGC) | 0.24(0.01) | 1.50 | 1.05(0.01) |
| BD | BD(PGC) | 0.18(0.01) | 1.27 | 1.02(0.01) |
| Years-of-schooling | EA(SSGAC) | 0.11(0.003) | 2.10 | 1.03(0.01) |
| General intelligence | GI(CTG) | 0.18(0.006) | 1.75 | 1.08(0.01) |

Abbreviations: ADHD, Attention-Deficit/Hyperactivity Disorder; ASD, Autism Spectrum Disorder; BD, Bipolar Disorder; EA; educational attainment; CTG, Complex Trait Genetics lab; GI, general intelligence; iPSYCH, The Lundbeck Foundation Initiative for Integrative Psychiatric Research; MDD, Major Depressive Disorder; PGC, Psychiatric Genomics Consortium; SCZ, Schizophrenia; SSGAC, Social Science Genetic Association Consortium; $\lambda_{GC}$, lambda GC; woADHD, without ADHD

SNP-heritability (SNP-$h^2$) was estimated with LDSC regression analysis[2]. SNP-$h^2$ estimates for EA and GI were calculated on the observed scale and for psychiatric disorders on a liability scale assuming a population prevalence of 0.012 (ASD), 0.05 (ADHD), 0.162 (MDD), 0.007 (SCZ) and 0.006 (BD).

Supplementary Table 3: Genetic correlations among psychiatric disorders

| Sample 1 | Sample 2 | $r_g$ (SE) | *P* |
|---|---|---|---|
| ASD(iPSYCH, woADHD) | ASD(PGC) | 0.84(0.11) | $<1x10^{-10}$ |
| | ADHD(iPSYCH) | 0.30 (0.06) | $2x10^{-7}$ |
| | MDD(PGC) | 0.43 (0.05) | $<1x10^{-10}$ |
| | SCZ(PGC) | 0.21 (0.06) | $1x10^{-4}$ |
| | BD(PGC) | 0.18 (0.06) | 0.001 |
| ASD(PGC) | ADHD(iPSYCH) | 0.04(0.08) | 0.61 |
| | MDD(PGC) | 0.12(0.05) | 0.017 |
| | SCZ(PGC) | 0.20(0.06) | 0.001 |
| | BD(PGC) | 0.12(0.06) | 0.063 |
| ADHD(iPSYCH) | MDD(PGC) | 0.55 (0.04) | $<1x10^{-10}$ |
| | SCZ(PGC) | 0.12 (0.04) | 0.004 |
| | BD(PGC) | 0.12 (0.05) | 0.007 |

Abbreviations: ADHD, Attention-Deficit/Hyperactivity Disorder; ASD, Autism Spectrum Disorder; BD, Bipolar Disorder; iPSYCH, The Lundbeck Foundation Initiative for Integrative Psychiatric Research; MDD; Major Depressive Disorder; PGC, Psychiatric Genomics Consortium; $r_g$, genetic correlation; SCZ, Schizophrenia; woADHD, without ADHD

Genetic correlations ($r_g$) among psychiatric disorder samples were estimated using summary statistics and unconstrained LD score correlation[3].

Supplementary Table 4: Genetic correlations of psychiatric disorders with educational attainment

| Sample 1 | Sample 2 | $r_g$ (SE) | *P* |
|---|---|---|---|
| EA (SSGAC) | ASD(iPSYCH) | 0.16(0.03) | $4x10^{-7}$ |
| | ASD(iPSYCH, woADHD) | 0.23(0.03) | $<1x10^{-10}$ |
| | ASD(PGC) | 0.28(0.03) | $<1x10^{-10}$ |
| | ADHD(iPSYCH) | -0.49(0.03) | $<1x10^{-10}$ |
| | ASD(PGC)+ADHD(iPSYCH) | -0.33(0.03) | $<1x10^{-10}$ |
| | MDD(PGC) | -0.22 (0.03) | $<1x10^{-10}$ |
| | SCZ(PGC) | 0.07 (0.02) | 0.003 |
| | BD(PGC) | 0.18 (0.02) | $<1x10^{-10}$ |

Abbreviations: ADHD, Attention-Deficit/Hyperactivity Disorder; ASD, Autism Spectrum Disorder; BD, Bipolar Disorder; EA; educational attainment; iPSYCH, The Lundbeck Foundation Initiative for Integrative Psychiatric Research; MDD, Major Depressive Disorder; PGC, Psychiatric Genomics Consortium; $r_g$, genetic correlation; SCZ, Schizophrenia; SSGAC, Social Science Genetic Association Consortium; woADHD, without ADHD;

Genetic correlations ($r_g$) of psychiatric disorder samples with educational attainment (EA) were estimated using summary statistics for EA, ASD(iPSYCH), ASD(iPSYCH, woADHD), ASD(PGC), ADHD(iPSYCH), ASD(PGC)+ADHD(iPSYCH), MDD(PGC), SCZ(PGC) and BD(PGC), respectively. ASD(PGC)+ADHD(iPSYCH) summary statistics were created by performing a random-effect meta-analysis combining ASD(PGC) and ADHD(iPSYCH) summary statistics. $r_g$ estimates were calculated using unconstrained LD score correlation[3].

Supplementary Table 5: Genetic correlations of ASD and ADHD with general intelligence

| Sample 1 | Sample 2 | $r_g$ (SE) | *P* |
|---|---|---|---|
| GI(CTG) | ASD(iPSYCH) | 0.20(0.04) | $1x10^{-8}$ |
| | ASD(iPSYCH, woADHD) | 0.25(0.04) | $<1x10^{-10}$ |
| | ASD(PGC) | 0.19(0.04) | $2x10^{-7}$ |
| | ADHD(iPSYCH) | -0.33(0.03) | $<1x10^{-10}$ |
| | ASD(PGC)+ADHD(iPSYCH) | -0.22(0.03) | $<1x10^{-10}$ |

Abbreviations: ADHD, Attention-Deficit/Hyperactivity Disorder; ASD, Autism Spectrum Disorder; CTG, Complex Trait Genetics lab; GI, general intelligence; iPSYCH, The Lundbeck Foundation Initiative for Integrative Psychiatric Research; PGC, Psychiatric Genomics Consortium; rg, genetic correlation; woADHD, without ADHD;

Genetic correlations ($r_g$) of ASD and ADHD samples with GI(CTG) were estimated using summary statistics for GI(CTG), ASD(iPSYCH), ASD(iPSYCH, woADHD), ASD(PGC), ADHD(iPSYCH) and ASD(PGC)+ADHD(iPSYCH), respectively. Latter were created by performing a random-effect meta-analysis combining ASD(PGC) and ADHD(iPSYCH) summary statistics. $r_g$ estimates were calculated using unconstrained LD score correlation[3].

Supplementary Table 6: Discovery ASD-MVR ($5x10^{-8}<P_{thr}<0.5$)

| Variant selection | | | Cross-disorder | Intercept | | ASD-specific effects | | Cross-disorder ADHD effects | |
|---|---|---|---|---|---|---|---|---|---|
| Variants | $N_{SNPs}$ | $P_{thr}$ | | $\beta_{int}$(SE) | *P* | MVR $\beta_{ASD}$(SE) | *P* | MVR $\beta_{\otimes ADHD}$(SE) | *P* |
| ASD (iPSYCH, woADHD) | 3* | $5x10^{-8}$ | ADHD (iPSYCH) | NA | NA | NA | NA | NA | NA |
| | 5 | $5x10^{-7}$ | | -0.010(0.007) | 0.28 | 0.10(0.065) | 0.26 | -0.015(0.028) | 0.65 |
| | 26 | $5x10^{-6}$ | | 0.004(0.003) | 0.16 | -0.003(0.030) | 0.93 | -0.022(0.024) | 0.38 |
| | 121 | $5x10^{-5}$ | | 0.001(0.001) | 0.38 | 0.010(0.013) | 0.44 | -0.018(0.014) | 0.22 |
| | 816 | 0.0005 | | $2x10^{-4}(4x10^{-4})$ | 0.54 | 0.015(0.005) | 0.002 | -0.064(0.006) | $1x10^{-8}$ |
| | 1,973 | 0.0015 | | $0.001(2x10^{-4})$ | 0.008 | 0.009(0.003) | 0.002 | -0.029(0.004) | $<1x10^{-10}$ |
| | 5,399 | 0.005 | | $5x10^{-4}(1x10^{-4})$ | $3x10^{-4}$ | 0.010(0.002) | $6x10^{-9}$ | -0.028(0.002) | $<1x10^{-10}$ |
| | 35,921 | 0.05 | | $3x10^{-4}(4x10^{-5})$ | $<1x10^{-10}$ | 0.007(0.001) | $<1x10^{-10}$ | -0.022(0.001) | $<1x10^{-10}$ |
| | 62,589 | 0.1 | | $2x10^{-4}(3x10^{-5})$ | $2x10^{-10}$ | 0.007(0.001) | $<1x10^{-10}$ | -0.020(0.001) | $<1x10^{-10}$ |
| | 134,210 | 0.3 | | $1x-10^{-4}(2x10^{-5})$ | $3x10^{-9}$ | $0.007(4x10^{-4})$ | $<1x10^{-10}$ | $-0.018(4x10^{-4})$ | $<1x10^{-10}$ |
| | 185,632 | 0.5 | | $1x10^{-4}(2x10^{-5})$ | $<1x10^{-10}$ | $0.007(4x10^{-4})$ | $<1x10^{-10}$ | $-0.017(3x10^{-4})$ | $<1x10^{-10}$ |

Abbreviations: ADHD, Attention-Deficit/Hyperactivity Disorder; ASD, Autism Spectrum Disorder; iPSYCH, The Lundbeck Foundation Initiative for Integrative Psychiatric Research; $N_{SNPs}$, number of SNPs; $P_{thr}$, *P*-value threshold; woADHD, without ADHD

Sets of independent genetic variants were selected from ASD(iPSYCH, woADHD) GWAS statistics at different *P*-value thresholds. Corresponding SNP estimates for ASD, ADHD and EA were subsequently extracted from ASD(iPSYCH, woADHD), ADHD(iPSYCH) and EA(SSGAC) GWAS statistics, respectively. Unconstrained multivariable regressions (MVRs) were fitted to identify simultaneously disorder-specific and cross-disorder associations with EA using ASD variant sets. ASD-specific MVR associations with EA (MVR $\beta_{ASD}$) were estimated with ASD variant sets and corresponding ASD SNP estimates. ADHD cross-disorder effects (ASD-MVR $\beta_{\otimes ADHD}$) were assessed with ADHD SNP estimates for ASD variant sets. SNP effects for all genetic variants were aligned according to the risk-increasing allele for ASD. All MVR effects are presented with respect to years-of-schooling, i.e. per increase in log-odds of ASD or ADHD liability, respectively The multiple testing threshold is $P<0.0023$.

*MVR analyses for ASD-related variants ($P_{thr}<5x10^{-8}$) did not converge.

Supplementary Table 7: Discovery ADHD-MVR ($5x10^{-8}<P_{thr}<0.5$)

| Variant selection | | | Cross-disorder | Intercept | | ADHD-specific effects | | Cross-disorder ASD effects | |
|---|---|---|---|---|---|---|---|---|---|
| Variants | $N_{SNPs}$ | $P_{thr}$ | | $\beta_{int}$(SE) | *P* | MVR $\beta_{ASD}$(SE) | *P* | MVR $\beta_{\otimes ASD}$(SE) | *P* |
| ADHD (iPSYCH) | 10 | $5x10^{-8}$ | ASD (iPSYCH, woADHD) | -0.006(0.017) | 0.76 | -0.068(0.16) | 0.68 | 0.12(0.075) | 0.16 |
| | 22 | $5x10^{-7}$ | | -0.002(0.006) | 0.70 | -0.055(0.060) | 0.37 | 0.061(0.044) | 0.19 |
| | 61 | $5x10^{-6}$ | | -0.003(0.002) | 0.13 | -0.048(0.024) | 0.054 | 0.096(0.026) | $5x10^{-4}$ |
| | 242 | $5x10^{-5}$ | | -0.003(0.001) | 0.001 | -0.013(0.011) | 0.24 | 0.025(0.012) | 0.036 |
| | 1,202 | 0.0005 | | $-0.002(3x10^{-4})$ | $2x10^{-6}$ | -0.015(0.004) | 0.001 | 0.023(0.005) | $2x10^{-6}$ |
| | 2,717 | 0.0015 | | $-0.002(2x10^{-4})$ | $<1x10^{-10}$ | -0.012(0.003) | $4x10^{-5}$ | 0.022(0.003) | $<1x10^{-10}$ |
| | 6,781 | 0.005 | | $-0.001(1x10^{-4})$ | $<1x10^{-10}$ | -0.009(0.002) | $5x10^{-8}$ | 0.018(0.002) | $<1x10^{-10}$ |
| | 41,334 | 0.05 | | $-0.001(4x10^{-5})$ | $<1x10^{-10}$ | -0.009(0.001) | $<1x10^{-10}$ | 0.013(0.001) | $<1x10^{-10}$ |
| | 71,015 | 0.1 | | $-0.001(3x10^{-5})$ | $<1x10^{-10}$ | -0.009(0.001) | $<1x10^{-10}$ | 0.012(0.001) | $<1x10^{-10}$ |
| | 164,083 | 0.3 | | $-3x10^{-4}(2x10^{-5})$ | $<1x10^{-10}$ | $-0.010(4x10^{-4})$ | $<1x10^{-10}$ | $0.010(3x10^{-4})$ | $<1x10^{-10}$ |
| | 234,530 | 0.5 | | $-2x10^{-4}(1x10^{-5})$ | $<1x10^{-10}$ | $-0.011(4x10^{-4})$ | $<1x10^{-10}$ | $0.009(3x10^{-4})$ | $<1x10^{-10}$ |

Abbreviations: ADHD, Attention-Deficit/Hyperactivity Disorder; ASD, Autism Spectrum Disorder; iPSYCH, The Lundbeck Foundation Initiative for Integrative Psychiatric Research; $N_{SNPs}$, number of SNPs; $P_{thr}$, *P*-value threshold; woADHD, without ADHD

Sets of independent genetic variants were selected from ADHD(iPSYCH) GWAS statistics at different *P*-value thresholds. Corresponding SNP estimates for ASD, ADHD and EA were subsequently extracted from ASD(iPSYCH, woADHD), ADHD(iPSYCH) and EA(SSGAC) GWAS statistics, respectively. Unconstrained multivariable regressions (MVRs) were fitted to identify simultaneously disorder-specific and cross-disorder associations with EA using ADHD variant sets. ADHD-specific MVR associations with EA (MVR $\beta_{ADHD}$) were estimated with ADHD variant sets and corresponding ADHD SNP estimates. ASD cross-disorder effects (ADHD-MVR $\beta_{\otimes ASD}$) were assessed with ASD SNP estimates for ADHD variant sets. SNP effects for all genetic variants were aligned according to the risk-increasing allele for ADHD. All MVR effects are presented with respect to years-of-schooling, i.e. per increase in log-odds of ASD or ADHD liability, respectively. The multiple testing threshold is $P<0.0023$.

Supplementary Table 8: Discovery ASD-MVR and ADHD-MVR ($P_{thr}$<0.0015; $P_{thr}$<0.05)

| *ASD-MVR* | | | | | | | | | | | | |
|---|---|---|---|---|---|---|---|---|---|---|---|---|
| Variant selection | | | Cross-disorder | Intercept | | ASD-specific effects | | Cross-disorder ADHD effects | | VIF | Model fit compared to a single model | |
| Variants | $N_{SNPs}$ | $P_{thr}$ | | $\beta_{int}$(SE) | *P* | MVR $\beta_{ASD}$(SE) | *P* | MVR $\beta_{\otimes ADHD}$(SE) | *P* | | $\Delta R^2$ (%) | Δdeviance, *P* |
| ASD (iPSYCH, woADHD) | 1,973 | 0.0015 | ADHD (iPSYCH) | 0.001(2x10$^{-4}$) | 0.008 | -5x10$^{-6}$(0.003) | 1.00 | NA | | NA | NA | |
| | | | | 0.001(2x10$^{-4}$) | 0.008 | 0.009(0.003) | 0.002 | -0.029(0.004) | <1x10$^{-10}$ | 1.19 | 3.0 | 167.06, <1x10$^{-10}$ |
| | 35,921 | 0.05 | | 3x10$^{-4}$(4x10$^{-5}$) | 1x10$^{-9}$ | 0.001(0.001) | 0.17 | NA | NA | NA | NA | NA |
| | | | | 3x10$^{-4}$(4x10$^{-5}$) | <1x10$^{-10}$ | 0.007(0.001) | <1x10$^{-10}$ | -0.022(0.001) | <1x10$^{-10}$ | 1.11 | 2.1 | 1533.8, <1x10$^{-10}$ |
| *ADHD-MVR* | | | | | | | | | | | | |
| Variant selection | | | Cross-disorder | Intercept | | ADHD-specific effects | | Cross-disorder ASD effects | | VIF | Model fit compared to a single model | |
| Variants | $N_{SNPs}$ | $P_{thr}$ | | $\beta_{int}$(SE) | *P* | MVR $\beta_{ADHD}$(SE) | *P* | MVR $\beta_{\otimes ASD}$(SE) | *P* | | $\Delta R^2$ (%) | Δdeviance, *P* |
| ADHD (iPSYCH) | 2,717 | 0.0015 | ASD (iPSYCH, woADHD) | -0.001(2x10$^{-4}$) | <1x10$^{-10}$ | -0.004(0.003) | 0.16 | NA | | NA | NA | |
| | | | | -0.002(2x10$^{-4}$) | <1x10$^{-10}$ | -0.012(0.003) | 4x10$^{-5}$ | 0.022(0.003) | <1x10$^{-10}$ | 1.18 | 1.9 | 154.24, <1x10$^{-10}$ |
| | 41,334 | 0.05 | | -0.001(4x10$^{-5}$) | <1x10$^{-10}$ | -0.004(0.001) | <1x10$^{-10}$ | NA | | NA | NA | |
| | | | | -0.001(4x10$^{-5}$) | <1x10$^{-10}$ | -0.009(0.001) | <1x10$^{-10}$ | 0.013(0.001) | <1x10$^{-10}$ | 1.11 | 0.9 | 744.05, <1x10$^{-10}$ |

Abbreviations: ADHD, Attention-Deficit/Hyperactivity Disorder; ASD, Autism Spectrum Disorder; iPSYCH, The Lundbeck Foundation Initiative for Integrative Psychiatric Research; $N_{SNPs}$, number of SNPs; $P_{thr}$, *P*-value threshold; VIF, variance inflation factor; woADHD, without ADHD

Sets of independent genetic variants were selected from ASD(iPSYCH, woADHD) and ADHD(iPSYCH) GWAS statistics at different *P*-value thresholds ($P_{thr}$<0.0015, $P_{thr}$<0.05). Corresponding SNP estimates for ASD, ADHD and EA were subsequently extracted from ASD(iPSYCH, woADHD), ADHD(iPSYCH) and EA(SSGAC) GWAS statistics, respectively. Unconstrained multivariable regressions (MVRs) were fitted to identify simultaneously disorder-specific and cross-disorder associations with EA, using either ASD or ADHD variant sets. ASD-specific MVR associations with EA (MVR $\beta_{ASD}$) were estimated with ASD variant sets and corresponding ASD SNP estimates. ADHD-specific MVR associations with EA (MVR $\beta_{ADHD}$) were estimated with ADHD variant sets and corresponding ADHD SNP estimates. MVR cross-disorder effects were either assessed with ADHD SNP estimates for ASD variant sets (ASD-MVR $\beta_{\otimes ADHD}$), or ASD SNP estimates for ADHD variant sets (ADHD-MVR $\beta_{\otimes ASD}$). SNP effects in ASD-MVR models were aligned according to ASD risk, and SNP effects in ADHD-MVR according to ADHD risk. All effects are presented with respect to years-of-schooling, i.e. per increase in log-odds of ASD or ADHD liability, respectively. The multiple testing threshold is *P*<0.0023. The model fit of MVRs and univariable regressions was compared with likelihood-ratio tests.

Supplementary Table 9: Follow-up ASD-MVR and ADHD-MVR ($P_{thr}$<0.0015; $P_{thr}$<0.05), concordant variants

*ASD-MVR*

| Variant selection | | | Cross-disorder | Intercept | | ASD-specific effects | | Cross-disorder ADHD effects | | VIF | Model fit compared to a single model | |
|---|---|---|---|---|---|---|---|---|---|---|---|---|
| Concordant variants | $N_{SNPs}$ | $P_{thr}$ | | $\beta_{int}$(SE) | *P* | MVR $\beta_{ASD}$(SE) | *P* | MVR $\beta_{\otimes ADHD}$(SE) | *P* | | $\Delta R^2$ (%) | Δdeviance, *P* |
| ASD (iPSYCH, woADHD) | 1,716 | 0.0015 | ADHD (iPSYCH) | $4x10^{-4}(2x10^{-4})$ | 0.15 | 0.001(0.003) | 0.76 | NA | | NA | NA | |
| | | | | $4x10^{-4}(2x10^{-4})$ | 0.12 | 0.011(0.003) | 0.001 | -0.026(0.005) | $5x10^{-8}$ | 1.40 | 1.7 | 79.67, $4x10^{-8}$ |
| | 28,086 | 0.05 | | $3x10^{-5}(5x10^{-5})$ | 0.49 | 0.001(0.001) | 0.11 | NA | | NA | NA | |
| | | | | $1x10^{-4}(5x10^{-5})$ | 0.054 | 0.009(0.001) | $<1x10^{-10}$ | -0.020(0.001) | $<1x10^{-10}$ | 1.36 | 1.1 | 579.37, $<1x10^{-10}$ |

*ADHD-MVR*

| Variant selection | | | Cross-disorder | Intercept | | ADHD-specific effects | | Cross-disorder ASD effects | | VIF | Model fit compared to a single model | |
|---|---|---|---|---|---|---|---|---|---|---|---|---|
| Concordant variants | $N_{SNPs}$ | $P_{thr}$ | | $\beta_{int}$(SE) | *P* | MVR $\beta_{ADHD}$(SE) | *P* | MVR $\beta_{\otimes ASD}$(SE) | *P* | | $\Delta R^2$ (%) | Δdeviance, *P* |
| ADHD (iPSYCH) | 2,382 | 0.0015 | ASD (iPSYCH, woADHD) | $-0.001(2x10^{-4})$ | $8x10^{-10}$ | -0.004(0.003) | 0.16 | NA | | NA | NA | |
| | | | | $-0.001(2x10^{-4})$ | $2x10^{-10}$ | -0.013(0.003) | $4x10^{-5}$ | 0.022(0.004) | $1x10^{-8}$ | 1.36 | 1.3 | 92.59, $1x10^{-8}$ |
| | 32,176 | 0.05 | | $-4x10^{-4}(5x10^{-5})$ | $<1x10^{-10}$ | -0.005(0.001) | $3x10^{-10}$ | NA | | NA | NA | |
| | | | | $-0.001(5x10^{-5})$ | $<1x10^{-10}$ | -0.011(0.001) | $<1x10^{-10}$ | 0.013(0.001) | $<1x10^{-10}$ | 1.36 | 0.5 | 339.16, $<1x10^{-10}$ |

Abbreviations: ADHD, Attention-Deficit/Hyperactivity Disorder; ASD, Autism Spectrum Disorder; iPSYCH, The Lundbeck Foundation Initiative for Integrative Psychiatric Research; $N_{SNPs}$, number of SNPs; $P_{thr}$, *P*-value threshold; VIF, variance inflation factor; woADHD, without ADHD

Sets of independent genetic variants were selected from ASD(iPSYCH, woADHD) and ADHD(iPSYCH) GWAS statistics at different *P*-value thresholds ($P_{thr}$<0.0015, $P_{thr}$<0.05). Corresponding SNP estimates for ASD, ADHD and EA were subsequently extracted from ASD(iPSYCH, woADHD), ADHD(iPSYCH) and EA(SSGAC) GWAS statistics, respectively. Unconstrained multivariable regressions (MVRs) were fitted to identify simultaneously disorder-specific and cross-disorder associations with EA, using either ASD or ADHD variant sets. ASD-specific MVR associations with EA (MVR $\beta_{ASD}$) were estimated with ASD variant sets and corresponding ASD SNP estimates. ADHD-specific MVR associations with EA (MVR $\beta_{ADHD}$) were estimated with ADHD variant sets and corresponding ADHD SNP estimates. MVR cross-disorder effects were either assessed with ADHD SNP estimates for ASD variant sets (ASD-MVR $\beta_{\otimes ADHD}$) or ASD SNP estimates for ADHD variant sets (ADHD-MVR $\beta_{\otimes ASD}$). Only variants with concordant ASD and ADHD association effect directions were included in the model and SNP effects were aligned to increase risk for both disorders. All MVR effects are presented with respect to years-of-schooling, i.e. per increase in log-odds of ASD or ADHD liability, respectively. The multiple testing threshold is *P*<0.0023.The model fit of MVRs and univariable regressions was compared with likelihood-ratio tests.

Supplementary Table 10: Follow-up ASD-MVR and ADHD-MVR ($P_{thr}$<0.0015; $P_{thr}$<0.05), analyses with ASD(PGC) SNP estimates (predictor)

*ASD-MVR*

| Variant selection: Variants | $N_{SNPs}$ | $P_{thr}$ | Cross-disorder | Intercept: $β_{int}$(SE) | *P* | ASD-specific effects: MVR $β_{ASD}$(SE) | *P* | Cross-disorder ADHD effects: MVR $β_{⊗ADHD}$(SE) | *P* | VIF | Model fit compared to a single model: $ΔR^2$ (%) | Δdeviance, *P* |
|---|---|---|---|---|---|---|---|---|---|---|---|---|
| ASD (iPSYCH, ASDwoADHD) variants with ASD(PGC) estimates | 1,886 | 0.0015 | ADHD (iPSYCH) | -6x10$^{-5}$(1x10$^{-4}$) | 0.69 | 0.010(0.003) | 0.004 | NA | | NA | NA | |
| | | | | -5x10$^{-5}$(1x10$^{-4}$) | 0.70 | 0.010(0.003) | 0.003 | -0.003(0.003) | 0.31 | 1.00 | 0.05 | 2.98, 0.31 |
| | 33,845 | 0.05 | | -1x10$^{-5}$(3x10$^{-5}$) | 0.61 | 0.005(0.001) | <1x10$^{-10}$ | NA | | NA | NA | |
| | | | | -9x10$^{-6}$(3x10$^{-5}$) | 0.76 | 0.005(0.001) | <1x10$^{-10}$ | -0.007(0.001) | <1x10$^{-10}$ | 1.00 | 0.41 | 285.63, <1x10$^{-10}$ |

*ADHD-MVR*

| Variant selection: Variants | $N_{SNPs}$ | $P_{thr}$ | Cross-disorder | Intercept: $β_{int}$(SE) | *P* | ADHD-specific effects: MVR $β_{ADHD}$(SE) | *P* | Cross-disorder ASD effects: MVR $β_{⊗ASD}$(SE) | *P* | VIF | Model fit compared to a single model: $ΔR^2$ (%) | Δdeviance, *P* |
|---|---|---|---|---|---|---|---|---|---|---|---|---|
| ADHD (iPSYCH) | 2,627 | 0.0015 | ASD (PGC) | -0.001(2x10$^{-4}$) | <1x10$^{-10}$ | -0.004(0.003) | 0.14 | NA | | NA | NA | |
| | | | | -0.001(2x10$^{-4}$) | <1x10$^{-10}$ | -0.004(0.003) | 0.13 | 0.002(0.002) | 0.35 | 1.00 | 0.03 | 2.62, 0.35 |
| | 38,656 | 0.05 | | -0.001(4x10$^{-5}$) | <1x10$^{-10}$ | -0.005(0.001) | <1x10$^{-10}$ | NA | | NA | NA | |
| | | | | -0.001(4x10$^{-5}$) | <1x10$^{-10}$ | -0.005(0.001) | <1x10$^{-10}$ | 0.003(4x10$^{-4}$) | <1x10$^{-10}$ | 1.00 | 0.16 | 123.59, <1x10$^{-10}$ |

Abbreviations: ADHD, Attention-Deficit/Hyperactivity Disorder; ASD, Autism Spectrum Disorder; iPSYCH, The Lundbeck Foundation Initiative for Integrative Psychiatric Research; PGC, Psychiatric Genomics Consortium; $N_{SNPs}$, number of SNPs; $P_{thr}$, *P*-value threshold; VIF, variance inflation factor

Sets of independent genetic variants were selected from ASD(iPSYCH, woADHD) and ADHD(iPSYCH) GWAS statistics at different *P*-value thresholds ($P_{thr}$<0.0015, $P_{thr}$<0.05). Corresponding SNP estimates for ASD, ADHD and EA were subsequently extracted from ASD(PGC), ADHD(iPSYCH) and EA(SSGAC) GWAS statistics, respectively. Unconstrained multivariable regressions (MVRs) were fitted to identify simultaneously disorder-specific and cross-disorder associations with EA, using either ASD or ADHD variant sets. ASD-specific MVR associations with EA (MVR $β_{ASD}$) were estimated with ASD variant sets and corresponding ASD SNP estimates. ADHD-specific MVR associations with EA (MVR $β_{ADHD}$) were estimated with ADHD variant sets and corresponding ADHD SNP estimates. MVR cross-disorder effects were either assessed with ADHD SNP estimates for ASD variant sets (ASD-MVR $β_{⊗ADHD}$) or ASD SNP estimates for ADHD variant sets (ADHD-MVR $β_{⊗ASD}$). SNP effects for all genetic variants were aligned according to the risk-increasing allele for the disorder used for variant set selection. All MVR effects are presented with respect to years-of-schooling, i.e. per increase in log-odds of ASD or ADHD liability, respectively. The multiple testing threshold is *P*<0.0125. The model fit of MVRs and univariable regressions was compared with likelihood-ratio tests.

Supplementary Table 11: Follow-up ASD-MVR and ADHD-MVR ($P_{thr}$<0.0015; $P_{thr}$<0.05), analyses with general intelligence SNP estimates (outcome)

*ASD-MVR*

| Variant selection | | | Cross-disorder | Intercept | | ASD-specific effects | | Cross-disorder ADHD effects | | VIF | Model fit compared to a single model | |
|---|---|---|---|---|---|---|---|---|---|---|---|---|
| Variants | $N_{SNPs}$ | $P_{thr}$ | | $\beta_{int}$(SE) | *P* | MVR $\beta_{ASD}$(SE) | *P* | MVR $\beta_{\otimes ADHD}$(SE) | *P* | | $\Delta R^2$ (%) | Δdeviance, *P* |
| ASD (iPSYCH, woADHD) | 1,904 | 0.0015 | ADHD (iPSYCH) | 0.001(3x$10^{-4}$) | 0.01 | -0.002(0.004) | 0.59 | NA | | NA | NA | |
| | | | | 0.001(3x$10^{-4}$) | 0.008 | 0.008(0.004) | 0.05 | -0.033(0.005) | 5x$10^{-10}$ | 1.18 | 2.02 | 79.80, 4x$10^{-10}$ |
| | 34,626 | 0.05 | | 4x$10^{-4}$(7x$10^{-5}$) | 5x$10^{-9}$ | 5x$10^{-5}$(0.001) | 0.96 | NA | | NA | NA | |
| | | | | 4x$10^{-4}$(7x$10^{-5}$) | 5x$10^{-10}$ | 0.006(0.001) | 2x$10^{-8}$ | -0.021(0.001) | <1x$10^{-10}$ | 1.11 | 0.91 | 509.13, <1x$10^{-10}$ |

*ADHD-MVR*

| Variant selection | | | Cross-disorder | Intercept | | ADHD-specific effects | | Cross-disorder ASD effects | | VIF | Model fit compared to a single model | |
|---|---|---|---|---|---|---|---|---|---|---|---|---|
| Variants | $N_{SNPs}$ | $P_{thr}$ | | $\beta_{int}$(SE) | *P* | MVR $\beta_{ADHD}$(SE) | *P* | MVR $\beta_{\otimes ASD}$(SE) | *P* | | $\Delta R^2$ (%) | Δdeviance, *P* |
| ADHD (iPSYCH) | 2,614 | 0.0015 | ASD (iPSYCH, woADHD) | -0.001(3x$10^{-4}$) | 0.001 | -0.006(0.004) | 0.09 | NA | | NA | NA | |
| | | | | -0.001(3x$10^{-4}$) | 0.001 | -0.012(0.004) | 0.002 | 0.016(0.004) | 2x$10^{-4}$ | 1.18 | 0.55 | 29.57, 1x$10^{-4}$ |
| | 39,767 | 0.05 | | -6x$10^{-4}$(6x$10^{-5}$) | <1x$10^{-10}$ | -0.003 (0.001) | 0.003 | NA | | NA | NA | |
| | | | | -0.001(6x$10^{-5}$) | <1x$10^{-10}$ | -0.007(0.001) | 3x$10^{-10}$ | 0.012(0.001) | <1x$10^{-10}$ | 1.10 | 0.33 | 211.34, <1x$10^{-10}$ |

Abbreviations: ADHD, Attention-Deficit/Hyperactivity Disorder; ASD, Autism Spectrum Disorder; iPSYCH, The Lundbeck Foundation Initiative for Integrative Psychiatric Research; $N_{SNPs}$, number of SNPs; $P_{thr}$, *P*-value threshold; woADHD, without ADHD

Sets of independent genetic variants were selected from ASD(iPSYCH, woADHD) and ADHD(iPSYCH) GWAS statistics at different *P*-value thresholds ($P_{thr}$<0.0015, $P_{thr}$<0.05). Corresponding SNP estimates for ASD, ADHD and general intelligence were subsequently extracted from ASD(iPSYCH, woADHD) and ADHD(iPSYCH) and general intelligence (CTG lab) GWAS statistics, respectively. Unconstrained multivariable regressions (MVRs) were fitted to identify simultaneously disorder-specific and cross-disorder associations with general intelligence, using either ASD or ADHD variant sets. ASD-specific MVR associations with general intelligence (MVR $\beta_{ASD}$) were estimated with ASD variant sets and corresponding ASD SNP estimates. ADHD-specific MVR associations with general intelligence (MVR $\beta_{ADHD}$) were estimated with ADHD variant sets and corresponding ADHD SNP estimates. MVR cross-disorder effects were either assessed with ADHD SNP estimates for ASD variant sets (ASD-MVR $\beta_{\otimes ADHD}$) or ASD SNP estimates for ADHD variant sets (ADHD-MVR $\beta_{\otimes ASD}$). SNP effects for all variants were aligned according to the risk-increasing allele for the disorder used for instrument selection. All MVR effects are presented as standard deviation-related differences in general intelligence, i.e. per increase in log-odds of ASD or ADHD liability, respectively. The multiple testing threshold is *P*<0.0125.

Supplementary Table 12: Follow-up ASD-MVR ($P_{thr}$ ASD<0.0015 and $5x10^{-8}$< $P_{thr}$ ADHD < 1), analyses with variants meeting joint ASD and ADHD selection criteria

| Variant selection | | | | Intercept | | ASD-specific effects | | ADHD cross-disorder effects | |
|---|---|---|---|---|---|---|---|---|---|
| $P_{thr}$ ASD (iPSYCH, woADHD) | $N_{SNPs}$ | % of initial variants | $P_{thr}$ ADHD (iPSYCH) | $\beta_{int}$(SE) | *P* | MVR $\beta_{ASD}$(SE) | *P* | MVR $\beta_{\otimes ADHD}$(SE) | *P* |
| 0.0015 | 1 | 0.05 | $5x10^{-8}$ | NA | NA | NA | NA | NA | NA |
| | 3 | 0.15 | $5x10^{-7}$ | NA | NA | NA | NA | NA | NA |
| | 4 | 0.20 | $5x10^{-6}$ | -0.11(0.003) | 0.16 | 0.37(0.075) | 0.13 | -0.29(0.060) | 0.13 |
| | 13 | 0.66 | $5x10^{-5}$ | -0.009(0.005) | 0.11 | 0.24(0.15) | 0.14 | -0.18(0.14) | 0.24 |
| | 52 | 2.64 | 0.0005 | -0.004(0.002) | 0.04 | 0.16(0.047) | 0.001 | -0.16(0.048) | 0.002 |
| | 83 | 4.21 | 0.0015 | -0.003(0.001) | 0.010 | 0.15(0.025) | $1x10^{-7}$ | -0.15(0.025) | $1x10^{-7}$ |
| | 134 | 6.79 | 0.005 | -0.002(0.001) | 0.15 | 0.12(0.022) | $6x10^{-7}$ | -0.13(0.024) | $2x10^{-7}$ |
| | 393 | 19.92 | 0.05 | -0.001(0.001) | 0.28 | 0.045(0.009) | $4x10^{-7}$ | -0.063(0.011) | $7x10^{-9}$ |
| | 536 | 27.17 | 0.1 | $-4x10^{-4}(5x10^{-4})$ | 0.40 | 0.040(0.007) | $2x10^{-8}$ | -0.059(0.009) | $<1x10^{-10}$ |
| | 805 | 40.80 | 0.3 | $5x10^{-4}(4x10^{-4})$ | 0.21 | 0.021(0.005) | $4x10^{-5}$ | -0.044(0.006) | $<1x10^{-10}$ |
| | 914 | 46.33 | 0.5 | $0.001(3x10^{-4})$ | 0.12 | 0.017(0.004) | $1x10^{-4}$ | -0.039(0.006) | $<1x10^{-10}$ |
| | 1973 | 100 | 1 | $0.001(2x10^{-4})$ | 0.008 | 0.009(0.003) | 0.002 | -0.029(0.004) | $<1x10^{-10}$ |

Abbreviations: ADHD, Attention-Deficit/Hyperactivity Disorder; ASD, Autism Spectrum Disorder; iPSYCH, The Lundbeck Foundation Initiative for Integrative Psychiatric Research; MVR, multivariable regression; $N_{SNPs}$, number of SNPs; $P_{thr}$, *P*-value threshold; woADHD, without ADHD;

Sets of independent genetic variants were selected from ASD(iPSYCH, woADHD) and ADHD(iPSYCH) GWAS statistics at $P_{thr}$<0.0015. For ASD variants ($P_{thr}$ <0.0015), we identified SNPs that were also represented within the ADHD variant set across a range of thresholds ($5x10^{-8}\leq P_{thr}$ <0.5). ASD variants were identified as tagged, if a SNP within the ADHD variant set was within 500kb and LD-$r^2$≥0.6. Corresponding SNP estimates for ASD, ADHD and EA were subsequently extracted from ASD(iPSYCH, woADHD), ADHD(iPSYCH) and EA(SSGAC) GWAS statistics, respectively. Unconstrained multivariable regressions (MVRs) (Figure 1) were fitted to identify simultaneously ASD-specific and ADHD cross-disorder associations with EA using subsets of ASD variants, associated with both ASD and ADHD. ASD-specific MVR associations with EA (MVR $\beta_{ASD}$) were estimated with ASD SNP estimates. ADHD cross-disorder effects (ASD-MVR $\beta_{\otimes ADHD}$) were assessed with ADHD SNP estimates. SNP effects for all genetic variants were aligned according to the risk-increasing allele for ASD. All MVR effects are presented with respect to years-of-schooling, i.e. per increase in log-odds of ASD or ADHD liability, respectively. The multiple testing threshold is *P*<0.0011.

Supplementary Table 13: Follow-up ADHD-MVR ($P_{thr}$ ADHD<0.0015 and 5x10$^{-8}$<$P_{thr}$ ASD<1), analyses with variants meeting joint ASD and ADHD selection criteria

| Variant selection | | | | Intercept | | ADHD-specific effects | | ASD cross-disorder effects | |
|---|---|---|---|---|---|---|---|---|---|
| $P_{thr}$ ADHD (iPSYCH) | $N_{SNPs}$ | % of initial variants | $P_{thr}$ ASD (iPSYCH, woADHD) | $\beta_{int}$(SE) | *P* | MVR $\beta_{ADHD}$(SE) | *P* | MVR $\beta_{\otimes ASD}$(SE) | *P* |
| 0.0015 | 1 | 0.04 | 5x10$^{-8}$ | NA | NA | NA | NA | NA | NA |
| | 2 | 0.07 | 5x10$^{-7}$ | NA | NA | NA | NA | NA | NA |
| | 6 | 0.22 | 5x10$^{-6}$ | 0.003(0.002) | 0.23 | -0.17(0.046) | 0.033 | 0.11(0.032) | 0.042 |
| | 13 | 0.48 | 5x10$^{-5}$ | -0.003(0.002) | 0.17 | -0.076(0.073) | 0.32 | 0.098(0.058) | 0.12 |
| | 49 | 1.80 | 0.0005 | -0.003(0.002) | 0.060 | -0.15(0.043) | 0.001 | 0.16(0.037) | 1x10$^{-4}$ |
| | 83 | 3.05 | 0.0015 | -0.003(0.001) | 0.038 | -0.10(0.026) | 2x10$^{-4}$ | 0.12(0.022) | 9x10$^{-7}$ |
| | 145 | 5.34 | 0.005 | -0.002(0.001) | 0.050 | -0.070(0.022) | 0.002 | 0.086(0.019) | 1x10$^{-5}$ |
| | 473 | 17.41 | 0.05 | -0.001(0.001) | 0.044 | -0.026(0.010) | 0.010 | 0.032(0.010) | 0.001 |
| | 638 | 23.48 | 0.1 | -0.001(4x10$^{-4}$) | 0.047 | -0.023(0.008) | 0.003 | 0.026(0.008) | 0.001 |
| | 937 | 34.49 | 0.3 | -0.001(3x10$^{-4}$) | 0.003 | -0.018(0.005) | 5x10$^{-4}$ | 0.023(0.006) | 5x10$^{-5}$ |
| | 1020 | 37.54 | 0.5 | -0.001(3x10$^{-4}$) | 0.005 | -0.017(0.005) | 3x10$^{-4}$ | 0.018(0.005) | 3x10$^{-4}$ |
| | 2717 | 100 | 1 | -0.002(2x10$^{-4}$) | <1x10$^{-10}$ | -0.012(0.003) | 4x10$^{-5}$ | 0.022(0.003) | <1x10$^{-10}$ |

Abbreviations: ADHD, Attention-Deficit/Hyperactivity Disorder; ASD, Autism Spectrum Disorder; iPSYCH, The Lundbeck Foundation Initiative for Integrative Psychiatric Research; MVR, multivariable regression; $N_{SNPs}$, number of SNPs; $P_{thr}$, *P*-value threshold; woADHD, without ADHD;

Sets of independent genetic variants were selected from ASD(iPSYCH, woADHD) and ADHD(iPSYCH) GWAS statistics at $P_{thr}$<0.0015. For ADHD variants ($P_{thr}$<0.0015), we identified SNPs that were also represented within the ASD variant set across a range of thresholds (5x10$^{-8}$≤ $P_{thr}$ <0.5). ADHD variants were identified as tagged, if a SNP within the ASD variant set was within 500kb and LD-$r^2$≥0.6. Corresponding SNP estimates for ASD, ADHD and EA were subsequently extracted from ASD(iPSYCH, woADHD), ADHD(iPSYCH) and EA(SSGAC) GWAS statistics, respectively. Unconstrained multivariable regressions (MVRs) (Figure 1) were fitted to identify simultaneously ADHD-specific and ASD cross-disorder associations with EA using subsets of ADHD variants, associated with both ASD and ADHD. ADHD-specific MVR associations with EA (MVR $\beta_{ADHD}$) were estimated with ADHD SNP estimates. ASD cross-disorder effects (ADHD-MVR $\beta_{\otimes ASD}$) were assessed with ASD SNP estimates. SNP effects for all genetic variants were aligned according to the risk-increasing allele for ADHD. All MVR effects are presented with respect to years-of-schooling, i.e. per increase in log-odds of ASD or ADHD liability, respectively. The multiple testing threshold is *P*<0.0011

.

Supplementary Table 14: SNP estimates for ASD-MVR and ADHD-MVR variants sets meeting joint ASD and ADHD selection criteria ($P_{thr}$ ADHD<0.0015 and $P_{thr}$ ASD<0.0015)

| ASD-MVR: ASD variants tagged by ADHD variants | | | | | | | ADHD MVR: ADHD variants tagged by ASD variants | | | | | | | Mapped gene(s) |
|---|---|---|---|---|---|---|---|---|---|---|---|---|---|---|
| SNP | A1 | A2 | ASD(iPSYCH, woADHD) | | ADHD(iPSYCH) | | SNP | A1 | A2 | ASD(iPSYCH, woADHD) | | ADHD(iPSYCH) | | |
| | | | β(SE) | *P* | β(SE) | *P* | | | | β(SE) | *P* | β(SE) | *P* | |
| rs10478063 | T | G | 0.06(0.02) | 7x10$^{-4}$ | 0.06(0.02) | 8x10$^{-4}$ | rs6869021 | T | C | 0.06(0.02) | 1x10$^{-3}$ | 0.07(0.02) | 7x10$^{-5}$ | *STARD4-AS1* |
| rs10503223 | C | G | 0.10(0.03) | 4x10$^{-4}$ | 0.11(0.03) | 4x10$^{-4}$ | rs10503223 | C | G | 0.10(0.03) | 4x10$^{-4}$ | 0.11(0.03) | 4x10$^{-4}$ | *CSMD1* |
| rs1052607 | G | A | 0.11(0.03) | 7x10$^{-4}$ | 0.12(0.03) | 4x10$^{-5}$ | rs77881576 | A | G | 0.11(0.03) | 8x10$^{-4}$ | 0.13(0.03) | 2x10$^{-5}$ | *MAST2, LOC110117498-PIK3R, PIK3R3* |
| rs10764449 | C | T | 0.07(0.02) | 3x10$^{-4}$ | 0.06(0.02) | 1x10$^{-3}$ | rs10764449 | C | T | 0.07(0.02) | 3x10$^{-4}$ | 0.06(0.02) | 1x10$^{-3}$ | *KIAA1217* |
| rs10932543 | A | G | 0.09(0.02) | 4x10$^{-6}$ | 0.07(0.02) | 1x10$^{-4}$ | rs10932543 | A | G | 0.09(0.02) | 4x10$^{-6}$ | 0.07(0.02) | 1x10$^{-4}$ | *VWC2L* |
| rs11124404 | G | T | 0.21(0.06) | 6x10$^{-4}$ | 0.20(0.05) | 3x10$^{-4}$ | rs12328829 | T | C | 0.18(0.06) | 2x10$^{-3}$ | 0.21(0.05) | 4x10$^{-5}$ | |
| rs112635299 | T | G | 0.17(0.05) | 7x10$^{-4}$ | 0.15(0.05) | 1x10$^{-3}$ | rs112635299 | T | G | 0.17(0.05) | 7x10$^{-4}$ | 0.15(0.05) | 1x10$^{-3}$ | |
| rs1133878 | A | G | 0.06(0.02) | 1x10$^{-3}$ | 0.04(0.02) | 1x10$^{-2}$ | rs2623245 | T | A | 0.05(0.02) | 4x10$^{-3}$ | 0.05(0.02) | 1x10$^{-3}$ | *SCG3, TMOD3* |
| rs114068758 | T | C | 0.12(0.04) | 1x10$^{-3}$ | 0.09(0.03) | 4x10$^{-3}$ | rs150844750 | C | T | 0.10(0.04) | 6x10$^{-3}$ | 0.12(0.03) | 4x10$^{-4}$ | *EDAR, MIR4435-1, MIR4435-2, MIR4771-1, MIR4771-2, RANBP2* |
| rs114457163 | C | T | 0.30(0.07) | 4x10$^{-6}$ | 0.21(0.06) | 5x10$^{-4}$ | rs114457163 | C | T | 0.30(0.07) | 4x10$^{-6}$ | 0.21(0.06) | 5x10$^{-4}$ | |
| rs11656656 | A | G | 0.07(0.02) | 1x10$^{-4}$ | 0.06(0.02) | 3x10$^{-4}$ | rs8074682 | G | A | 0.07(0.02) | 1x10$^{-4}$ | 0.06(0.02) | 2x10$^{-4}$ | *MGAT5B* |
| rs116687082 | T | C | 0.16(0.05) | 4x10$^{-4}$ | 0.16(0.04) | 8x10$^{-5}$ | rs116687082 | T | C | 0.16(0.05) | 4x10$^{-4}$ | 0.16(0.04) | 8x10$^{-5}$ | |
| rs118058985 | T | C | 0.18(0.04) | 3x10$^{-6}$ | 0.14(0.03) | 5x10$^{-5}$ | rs118058985 | T | C | 0.18(0.04) | 3x10$^{-6}$ | 0.14(0.03) | 5x10$^{-5}$ | |
| rs1198850 | C | T | 0.08(0.02) | 3x10$^{-4}$ | 0.07(0.02) | 1x10$^{-3}$ | rs1198850 | C | T | 0.08(0.02) | 3x10$^{-4}$ | 0.07(0.02) | 1x10$^{-3}$ | *ATP6V1C2* |
| rs12142993 | G | A | 0.22(0.06) | 8x10$^{-5}$ | 0.15(0.05) | 1x10$^{-3}$ | rs12142993 | G | A | 0.22(0.06) | 8x10$^{-5}$ | 0.15(0.05) | 1x10$^{-3}$ | *FAM72C, FMO2, SRGAP2D* |
| rs12482859 | A | T | 0.10(0.03) | 1x10$^{-3}$ | 0.09(0.03) | 2x10$^{-3}$ | rs12482860 | C | T | 0.09(0.03) | 5x10$^{-3}$ | 0.09(0.03) | 1x10$^{-3}$ | |
| rs12900962 | C | T | 0.08(0.03) | 1x10$^{-3}$ | 0.07(0.02) | 3x10$^{-3}$ | rs3794565 | A | G | 0.08(0.03) | 2x10$^{-3}$ | 0.08(0.02) | 1x10$^{-3}$ | *MYO5A* |
| rs13207266 | A | C | 0.06(0.02) | 5x10$^{-4}$ | 0.05(0.02) | 2x10$^{-3}$ | rs9406227 | G | T | 0.06(0.02) | 1x10$^{-3}$ | 0.05(0.02) | 1x10$^{-3}$ | |
| rs13338042 | C | A | 0.18(0.04) | 4x10$^{-5}$ | 0.13(0.04) | 2x10$^{-3}$ | rs28422934 | G | C | 0.18(0.04) | 5x10$^{-5}$ | 0.13(0.04) | 8x10$^{-4}$ | |
| rs139028896 | G | C | 0.12(0.04) | 1x10$^{-3}$ | 0.11(0.03) | 7x10$^{-4}$ | rs41300124 | C | T | 0.11(0.04) | 4x10$^{-3}$ | 0.12(0.03) | 5x10$^{-4}$ | *CELA3A* |
| rs146374277 | C | T | 0.25(0.08) | 1x10$^{-3}$ | 0.23(0.07) | 7x10$^{-4}$ | rs146374277 | C | T | 0.25(0.08) | 1x10$^{-3}$ | 0.23(0.07) | 7x10$^{-4}$ | |
| rs148532090 | T | C | 0.29(0.07) | 3x10$^{-5}$ | 0.28(0.06) | 1x10$^{-5}$ | rs148532090 | T | C | 0.29(0.07) | 3x10$^{-5}$ | 0.28(0.06) | 1x10$^{-5}$ | *IP6K2* |
| rs1554380 | A | G | 0.10(0.03) | 2x10$^{-4}$ | 0.08(0.03) | 2x10$^{-3}$ | rs17012945 | G | A | 0.10(0.03) | 1x10$^{-3}$ | 0.09(0.03) | 4x10$^{-4}$ | |
| rs17477236 | C | A | 0.08(0.02) | 1x10$^{-4}$ | 0.06(0.02) | 1x10$^{-3}$ | rs17477236 | C | A | 0.08(0.02) | 1x10$^{-4}$ | 0.06(0.02) | 1x10$^{-3}$ | *VAV3* |
| rs184032510 | C | T | 0.20(0.06) | 6x10$^{-4}$ | 0.14(0.05) | 9x10$^{-3}$ | rs141735644 | T | C | 0.21(0.06) | 1x10$^{-3}$ | 0.21(0.06) | 4x10$^{-4}$ | |

| | | | | | | | | | | | | | | |
|---|---|---|---|---|---|---|---|---|---|---|---|---|---|---|
| rs1985169 | A | G | 0.11(0.02) | 2x10$^{-6}$ | 0.09(0.02) | 7x10$^{-6}$ | rs112337001 | T | C | 0.10(0.02) | 1x10$^{-6}$ | 0.08(0.02) | 7x10$^{-6}$ | |
| rs202823 | A | G | 0.16(0.04) | 6x10$^{-5}$ | 0.13(0.04) | 2x10$^{-4}$ | rs17664036 | C | T | 0.14(0.04) | 1x10$^{-4}$ | 0.12(0.03) | 1x10$^{-4}$ | *XRN2, KIZ* |
| rs2228048 | T | C | 0.21(0.06) | 1x10$^{-3}$ | 0.18(0.06) | 2x10$^{-3}$ | rs3773648 | C | G | 0.20(0.07) | 2x10$^{-3}$ | 0.19(0.06) | 1x10$^{-3}$ | *TGFBR2* |
| rs2239113 | G | A | 0.07(0.02) | 9x10$^{-4}$ | 0.07(0.02) | 2x10$^{-4}$ | rs4765959 | A | T | 0.06(0.02) | 2x10$^{-3}$ | 0.07(0.02) | 9x10$^{-5}$ | *CACNA1C* |
| rs2328878 | G | A | 0.06(0.02) | 1x10$^{-3}$ | 0.05(0.02) | 3x10$^{-3}$ | rs2744296 | A | G | 0.04(0.02) | 1x10$^{-2}$ | 0.05(0.02) | 6x10$^{-4}$ | *CARMIL1* |
| rs2391769 | G | A | 0.09(0.02) | 5x10$^{-7}$ | 0.09(0.02) | 8x10$^{-8}$ | rs2391769 | G | A | 0.09(0.02) | 5x10$^{-7}$ | 0.09(0.02) | 8x10$^{-8}$ | |
| rs2635182 | T | C | 0.07(0.02) | 6x10$^{-5}$ | 0.05(0.02) | 2x10$^{-3}$ | rs250274 | T | G | 0.06(0.02) | 2x10$^{-4}$ | 0.06(0.02) | 2x10$^{-4}$ | |
| rs33823 | T | C | 0.07(0.02) | 3x10$^{-5}$ | 0.04(0.02) | 6x10$^{-3}$ | rs57457691 | C | T | 0.05(0.02) | 1x10$^{-2}$ | 0.05(0.02) | 1x10$^{-3}$ | *PEPD* |
| rs35357088 | C | T | 0.13(0.03) | 2x10$^{-4}$ | 0.10(0.03) | 1x10$^{-3}$ | rs77529534 | C | T | 0.13(0.04) | 9x10$^{-4}$ | 0.13(0.03) | 2x10$^{-4}$ | *TEK* |
| rs36092443 | A | G | 0.15(0.04) | 4x10$^{-4}$ | 0.15(0.04) | 9x10$^{-5}$ | rs36092443 | A | G | 0.15(0.04) | 4x10$^{-4}$ | 0.15(0.04) | 9x10$^{-5}$ | |
| rs3767602 | G | A | 0.13(0.04) | 4x10$^{-4}$ | 0.11(0.03) | 8x10$^{-4}$ | rs11810580 | C | T | 0.11(0.04) | 3x10$^{-3}$ | 0.11(0.03) | 7x10$^{-4}$ | *TMIGD3* |
| rs416223 | C | A | 0.06(0.02) | 5x10$^{-4}$ | 0.07(0.02) | 2x10$^{-6}$ | rs325506 | C | G | 0.05(0.02) | 3x10$^{-3}$ | 0.08(0.02) | 5x10$^{-7}$ | |
| rs4322805 | A | G | 0.06(0.02) | 2x10$^{-4}$ | 0.06(0.02) | 3x10$^{-4}$ | rs4322805 | A | G | 0.06(0.02) | 2x10$^{-4}$ | 0.06(0.02) | 3x10$^{-4}$ | |
| rs4609618 | C | A | 0.08(0.02) | 9x10$^{-6}$ | 0.05(0.02) | 7x10$^{-4}$ | rs4609618 | C | A | 0.08(0.02) | 9x10$^{-6}$ | 0.05(0.02) | 7x10$^{-4}$ | |
| rs4679557 | A | G | 0.06(0.02) | 8x10$^{-4}$ | 0.05(0.02) | 9x10$^{-4}$ | rs6787365 | A | G | 0.06(0.02) | 1x10$^{-3}$ | 0.06(0.02) | 5x10$^{-4}$ | |
| rs4916723 | C | A | 0.07(0.02) | 1x10$^{-4}$ | 0.09(0.02) | 5x10$^{-9}$ | rs4916723 | C | A | 0.07(0.02) | 1x10$^{-4}$ | 0.09(0.02) | 5x10$^{-9}$ | *LINC00461* |
| rs4952312 | A | G | 0.10(0.03) | 5x10$^{-4}$ | 0.08(0.03) | 2x10$^{-3}$ | rs77384847 | C | T | 0.09(0.03) | 8x10$^{-4}$ | 0.08(0.03) | 1x10$^{-3}$ | *LINC00486* |
| rs4981706 | A | T | 0.07(0.02) | 6x10$^{-4}$ | 0.07(0.02) | 5x10$^{-4}$ | rs4981706 | A | T | 0.07(0.02) | 6x10$^{-4}$ | 0.07(0.02) | 5x10$^{-4}$ | |
| rs501371 | G | A | 0.18(0.05) | 1x10$^{-4}$ | 0.14(0.04) | 7x10$^{-4}$ | rs1277718 | C | G | 0.18(0.05) | 1x10$^{-4}$ | 0.14(0.04) | 7x10$^{-4}$ | |
| rs529507 | G | A | 0.14(0.02) | 3x10$^{-8}$ | 0.07(0.02) | 2x10$^{-3}$ | rs478324 | A | C | 0.13(0.02) | 1x10$^{-7}$ | 0.07(0.02) | 1x10$^{-3}$ | *NTM* |
| rs58689390 | C | T | 0.07(0.02) | 8x10$^{-4}$ | 0.04(0.02) | 2x10$^{-2}$ | rs10416388 | G | A | 0.07(0.02) | 1x10$^{-3}$ | 0.06(0.02) | 1x10$^{-3}$ | |
| rs59884341 | C | T | 0.22(0.06) | 5x10$^{-4}$ | 0.22(0.06) | 2x10$^{-4}$ | rs59884341 | C | T | 0.22(0.06) | 5x10$^{-4}$ | 0.22(0.06) | 2x10$^{-4}$ | |
| rs6029251 | G | A | 0.06(0.02) | 8x10$^{-4}$ | 0.07(0.02) | 8x10$^{-5}$ | rs735031 | G | A | 0.06(0.02) | 8x10$^{-4}$ | 0.07(0.02) | 8x10$^{-5}$ | |
| rs60798171 | G | T | 0.07(0.02) | 4x10$^{-4}$ | 0.07(0.02) | 6x10$^{-5}$ | rs60798171 | G | T | 0.07(0.02) | 4x10$^{-4}$ | 0.07(0.02) | 6x10$^{-5}$ | |
| rs62192813 | G | T | 0.18(0.06) | 1x10$^{-3}$ | 0.18(0.05) | 4x10$^{-4}$ | rs62192813 | G | T | 0.18(0.06) | 1x10$^{-3}$ | 0.18(0.05) | 4x10$^{-4}$ | |
| rs638311 | A | T | 0.07(0.02) | 1x10$^{-4}$ | 0.05(0.02) | 7x10$^{-4}$ | rs10802808 | T | C | 0.07(0.02) | 2x10$^{-4}$ | 0.06(0.02) | 6x10$^{-5}$ | *CHRM3* |
| rs6421157 | G | A | 0.07(0.02) | 2x10$^{-5}$ | 0.05(0.02) | 3x10$^{-4}$ | rs10454999 | G | A | 0.07(0.02) | 8x10$^{-5}$ | 0.06(0.02) | 9x10$^{-5}$ | |
| rs6422311 | G | A | 0.07(0.02) | 1x10$^{-4}$ | 0.07(0.02) | 2x10$^{-5}$ | rs6858688 | A | T | 0.07(0.02) | 2x10$^{-4}$ | 0.07(0.02) | 2x10$^{-5}$ | |
| rs6435661 | C | T | 0.07(0.02) | 2x10$^{-4}$ | 0.06(0.02) | 1x10$^{-3}$ | rs6435661 | C | T | 0.07(0.02) | 2x10$^{-4}$ | 0.06(0.02) | 1x10$^{-3}$ | *ERBB4* |
| rs6467754 | G | A | 0.08(0.02) | 4x10$^{-5}$ | 0.05(0.02) | 1x10$^{-3}$ | rs4732306 | C | T | 0.07(0.02) | 8x10$^{-5}$ | 0.06(0.02) | 8x10$^{-4}$ | |
| rs6584356 | A | C | 0.15(0.04) | 2x10$^{-4}$ | 0.13(0.04) | 5x10$^{-4}$ | rs35662118 | T | C | 0.15(0.04) | 2x10$^{-4}$ | 0.13(0.04) | 4x10$^{-4}$ | *PKD2L1* |
| rs6701243 | A | C | 0.08(0.02) | 2x10$^{-5}$ | 0.06(0.02) | 5x10$^{-4}$ | rs35518820 | C | T | 0.07(0.02) | 2x10$^{-4}$ | 0.06(0.02) | 4x10$^{-4}$ | |
| rs6750890 | T | G | 0.10(0.02) | 3x10$^{-5}$ | 0.07(0.02) | 8x10$^{-4}$ | rs6750890 | T | G | 0.10(0.02) | 3x10$^{-5}$ | 0.07(0.02) | 8x10$^{-4}$ | *LOC100507443* |
| rs71429234 | G | A | 0.07(0.02) | 1x10$^{-3}$ | 0.06(0.02) | 2x10$^{-3}$ | rs13006868 | G | A | 0.07(0.02) | 2x10$^{-3}$ | 0.06(0.02) | 1x10$^{-3}$ | |
| rs72660658 | C | T | 0.06(0.02) | 1x10$^{-3}$ | 0.06(0.02) | 2x10$^{-4}$ | rs10488866 | T | C | 0.06(0.02) | 1x10$^{-3}$ | 0.06(0.02) | 2x10$^{-4}$ | *CXXC4-AS1* |

| | | | | | | | | | | | | | | |
|---|---|---|---|---|---|---|---|---|---|---|---|---|---|---|
| rs72762883 | T | G | 0.11(0.03) | 3x10$^{-4}$ | 0.08(0.03) | 5x10$^{-3}$ | rs72762892 | A | G | 0.10(0.03) | 7x10$^{-4}$ | 0.08(0.03) | 8x10$^{-4}$ | *KIF26B* |
| rs73086963 | T | C | 0.25(0.06) | 1x10$^{-4}$ | 0.17(0.06) | 6x10$^{-3}$ | rs41308228 | A | T | 0.24(0.07) | 3x10$^{-4}$ | 0.21(0.06) | 7x10$^{-4}$ | *CACNA2D2* |
| rs73088112 | C | T | 0.27(0.07) | 1x10$^{-4}$ | 0.23(0.06) | 4x10$^{-4}$ | rs73074869 | A | G | 0.27(0.07) | 1x10$^{-4}$ | 0.24(0.06) | 2x10$^{-4}$ | *RHOA, BSN* |
| rs73116288 | G | T | 0.07(0.02) | 1x10$^{-4}$ | 0.06(0.02) | 7x10$^{-4}$ | rs62254854 | C | T | 0.07(0.02) | 3x10$^{-4}$ | 0.06(0.02) | 2x10$^{-4}$ | |
| rs7319835 | T | C | 0.09(0.03) | 1x10$^{-3}$ | 0.09(0.03) | 4x10$^{-4}$ | rs61965320 | A | G | 0.09(0.03) | 2x10$^{-3}$ | 0.09(0.03) | 4x10$^{-4}$ | |
| rs73214716 | T | C | 0.11(0.04) | 1x10$^{-3}$ | 0.11(0.03) | 4x10$^{-4}$ | rs73214716 | T | C | 0.11(0.04) | 1x10$^{-3}$ | 0.11(0.03) | 4x10$^{-4}$ | |
| rs74992427 | C | T | 0.20(0.06) | 1x10$^{-3}$ | 0.17(0.06) | 2x10$^{-3}$ | rs77709635 | T | C | 0.16(0.06) | 5x10$^{-3}$ | 0.20(0.05) | 9x10$^{-5}$ | *ZNF704* |
| rs75263467 | A | G | 0.16(0.04) | 3x10$^{-4}$ | 0.21(0.04) | 5x10$^{-7}$ | rs75263467 | A | G | 0.16(0.04) | 3x10$^{-4}$ | 0.21(0.04) | 5x10$^{-7}$ | |
| rs75874195 | A | G | 0.15(0.04) | 9x10$^{-4}$ | 0.10(0.04) | 1x10$^{-2}$ | rs78924818 | C | G | 0.17(0.06) | 3x10$^{-3}$ | 0.18(0.05) | 3x10$^{-4}$ | |
| rs7625233 | G | A | 0.06(0.02) | 4x10$^{-4}$ | 0.06(0.02) | 1x10$^{-4}$ | rs7629352 | G | A | 0.05(0.02) | 5x10$^{-3}$ | 0.07(0.02) | 4x10$^{-5}$ | |
| rs76530346 | G | A | 0.10(0.03) | 1x10$^{-3}$ | 0.09(0.03) | 1x10$^{-3}$ | rs78826784 | G | A | 0.09(0.03) | 2x10$^{-3}$ | 0.09(0.03) | 4x10$^{-4}$ | *AUH, LINC00484* |
| rs7781266 | C | A | 0.07(0.02) | 1x10$^{-3}$ | -0.05(0.02) | 1x10$^{-2}$ | rs17167170 | G | A | -0.06(0.02) | 8x10$^{-3}$ | 0.06(0.02) | 1x10$^{-3}$ | *EXOC4, LOC101928861* |
| rs77966298 | A | G | 0.09(0.02) | 2x10$^{-4}$ | 0.09(0.02) | 5x10$^{-5}$ | rs6737620 | C | T | 0.08(0.02) | 6x10$^{-4}$ | 0.09(0.02) | 1x10$^{-5}$ | |
| rs78491961 | T | C | 0.21(0.06) | 3x10$^{-4}$ | 0.22(0.06) | 3x10$^{-4}$ | rs78491961 | T | C | 0.21(0.06) | 3x10$^{-4}$ | 0.22(0.06) | 3x10$^{-4}$ | |
| rs78737628 | C | A | 0.09(0.02) | 4x10$^{-4}$ | 0.08(0.02) | 5x10$^{-4}$ | rs78737628 | C | A | 0.09(0.02) | 4x10$^{-4}$ | 0.08(0.02) | 5x10$^{-4}$ | |
| rs80013344 | A | G | 0.10(0.03) | 3x10$^{-4}$ | 0.09(0.02) | 3x10$^{-4}$ | rs4947694 | C | A | 0.10(0.03) | 3x10$^{-4}$ | 0.11(0.03) | 2x10$^{-5}$ | |
| rs80088989 | C | A | 0.15(0.05) | 1x10$^{-3}$ | 0.17(0.05) | 2x10$^{-4}$ | rs80088989 | C | A | 0.15(0.05) | 1x10$^{-3}$ | 0.17(0.05) | 2x10$^{-4}$ | *TEK* |
| rs80229434 | A | G | 0.22(0.07) | 1x10$^{-3}$ | 0.19(0.06) | 3x10$^{-3}$ | rs111620031 | C | G | 0.14(0.07) | 3x10$^{-2}$ | 0.20(0.06) | 1x10$^{-3}$ | *MIR3680-1, MIR3680-2* |
| rs927603 | T | C | 0.06(0.02) | 3x10$^{-4}$ | 0.04(0.02) | 6x10$^{-3}$ | rs8016878 | A | G | 0.05(0.02) | 5x10$^{-3}$ | 0.06(0.02) | 1x10$^{-4}$ | |
| rs978216 | C | T | 0.06(0.02) | 8x10$^{-4}$ | 0.06(0.02) | 2x10$^{-5}$ | rs978216 | C | T | 0.06(0.02) | 8x10$^{-4}$ | 0.06(0.02) | 2x10$^{-5}$ | |
| rs9816530 | T | G | 0.08(0.02) | 6x10$^{-4}$ | 0.06(0.02) | 4x10$^{-3}$ | rs6441815 | T | C | 0.07(0.02) | 1x10$^{-3}$ | 0.06(0.02) | 1x10$^{-3}$ | |
| rs9855048 | G | A | 0.12(0.03) | 1x10$^{-4}$ | 0.10(0.03) | 4x10$^{-4}$ | rs9855048 | G | A | 0.12(0.03) | 1x10$^{-4}$ | 0.10(0.03) | 4x10$^{-4}$ | |
| rs9878955 | G | A | 0.06(0.02) | 1x10$^{-3}$ | 0.04(0.02) | 1x10$^{-2}$ | rs7616968 | G | A | 0.05(0.02) | 1x10$^{-2}$ | 0.06(0.02) | 5x10$^{-4}$ | *OSTN* |

Abbreviations: ADHD, Attention-Deficit/Hyperactivity Disorder; ASD, Autism Spectrum Disorder; iPSYCH, The Lundbeck Foundation Initiative for Integrative Psychiatric Research; $P_{thr}$, *P*-value threshold; woADHD, without ADHD;

ASD(iPSYCH, woADHD) and ADHD(iPSYCH) SNP estimates for ASD variants ($P_{thr}$ <0.0015) that were also represented within the ADHD variant set ($P_{thr}$ <0.0015, 500kb and LD-$r^2$≥0.6). Likewise, SNP estimates for ADHD variants ($P_{thr}$ <0.0015) that were also represented within the ASD variant set ($P_{thr}$ <0.0015, 500kb and LD-$r^2$≥0.6) are shown. ASD and ADHD variants that tag each other are represented on the same row. All SNP estimates are aligned according to A1. Mapped genes were identified using a 0kb window range.

Supplementary Table 15: Permutation analysis

| MVR | Variant selection | | | Empirical *P*-value (SE) | |
|---|---|---|---|---|---|
| | $P_{thr}$ ASD (iPSYCH, woADHD) | $P_{thr}$ ADHD (iPSYCH) | $N_{SNPs}$ | Specific effects | Cross-disorder effects |
| **ASD-MVR** | 0.0015 | 0.0015 | 83 | $<1x10^{-4}$($<1x10^{-4}$) | $7x10^{-4}$($3x10^{-4}$) |
| **ADHD-MVR** | 0.0015 | 0.0015 | 83 | $9x10^{-4}$($3x10^{-4}$) | $2x10^{-4}$($1x10^{-4}$) |

Abbreviations: ADHD, Attention-Deficit/Hyperactivity Disorder; ASD, Autism Spectrum Disorder; iPSYCH, The Lundbeck Foundation Initiative for Integrative Psychiatric Research; MVR, multivariable regression; $N_{SNPs}$, number of SNPs; $P_{thr}$, *P*-value threshold; woADHD, without ADHD;

83 SNPs were randomly selected from either ASD variants ($P_{thr}$<0.0015) or ADHD variants ($P_{thr}$<0.0015). Corresponding SNP estimates for ASD, ADHD and EA were subsequently extracted from ASD(iPSYCH, woADHD), ADHD(iPSYCH) and EA(SSGAC) GWAS statistics, respectively. Unconstrained MVRs were performed and the number of times a permuted MVR effect was at least as significant as an observed MVR effect counted. The total number of permutations performed was 10,000.

Supplementary Table 16: Follow-up ASD-MVR ($P_{thr}$<0.0015; $P_{thr}$<0.05), analyses with MDD, SCZ and BD SNP estimates (predictor)

| Variant selection | | | Cross-disorder | Intercept | | ASD-specific effect | | Cross-disorder effect | | VIF | Model fit compared to a single model | |
|---|---|---|---|---|---|---|---|---|---|---|---|---|
| Variants | $N_{SNPs}$ | $P_{thr}$ | | $\beta_{int}$(SE) | *P* | MVR $\beta_{ASD}$(SE) | *P* | MVR $\beta_{\otimes}$(SE) | *P* | | $\Delta R^2$ (%) | Δdeviance, *P* |
| ASD (iPSYCH, woADHD) | 2,644 | 0.0015 | MDD (PGC) | 0.001(1x10$^{-4}$) | 3x10$^{-5}$ | -0.001(0.002) | 0.45 | NA | | NA | NA | |
| | | | | 0.001(2x10$^{-4}$) | 3x10$^{-5}$ | -0.001(0.002) | 0.55 | $\beta_{\otimes MDD}$=-0.004(0.006) | 0.46 | 1.04 | 0.02 | 1.37, 0.45 |
| | 50,904 | 0.05 | | 3x10$^{-4}$ (4x10$^{-5}$) | <1x10$^{-10}$ | 7x10$^{-4}$(5x10$^{-4}$) | 0.13 | NA | | NA | NA | |
| | | | | 3x10$^{-4}$(4x10$^{-5}$) | <1x10$^{-10}$ | 0.001(5x10$^{-4}$) | 0.002 | $\beta_{\otimes MDD}$=-0.012(0.001) | <1x10$^{-10}$ | 1.02 | 0.29 | 264.96, <1x10$^{-10}$ |
| | 1,786 | 0.0015 | SCZ (PGC) | 0.001(3x10$^{-4}$) | 0.01 | -0.001(0.003) | 0.78 | NA | | NA | NA | |
| | | | | 0.001(3x10$^{-4}$) | 0.02 | -0.001(0.003) | 0.83 | $\beta_{\otimes SCZ}$=0.016(0.004) | 3x10$^{-4}$ | 1.00 | 0.71 | 37.92, 3x10$^{-4}$ |
| | 31,026 | 0.05 | | 2x10$^{-4}$(5x10$^{-5}$ | 6x10$^{-5}$ | 0.002(8x10$^{-4}$) | 0.01 | NA | | NA | NA | |
| | | | | 2x10$^{-4}$(5x10$^{-5}$) | 1x10$^{-4}$ | 0.002(0.001) | 0.02 | $\beta_{\otimes SCZ}$ =0.006(0.001) | 3x10$^{-9}$ | 1.00 | 0.11 | 73.31, 3x10$^{-9}$ |
| | 1,859 | 0.0015 | BD (PGC) | 0.001(2x10$^{-4}$) | 0.003 | -0.001(0.003) | 0.61 | | | | | |
| | | | | 0.001(2x10$^{-4}$) | 0.004 | -0.001(0.003) | 0.61 | $\beta_{\otimes BD}$=0.019(0.004) | 6x10$^{-7}$ | 1.00 | 1.33 | 71.40, 6x10$^{-7}$ |
| | 32,367 | 0.05 | | 2x10$^{-4}$(5x10$^{-5}$) | 2x10$^{-6}$ | 0.002(7x10$^{-4}$) | 0.04 | | | | | |
| | | | | 2x10$^{-4}$(5x10$^{-5}$) | 7x10$^{-6}$ | 0.002(0.001) | 0.033 | $\beta_{\otimes BD}$ =0.009(0.001) | <1x10$^{-10}$ | 1.00 | 0.41 | 272.24, <1x10$^{-10}$ |

Abbreviations: ASD, Autism Spectrum Disorder; BD, Bipolar Disorder; iPSYCH, The Lundbeck Foundation Initiative for Integrative Psychiatric Research; MDD; Major Depressive Disorder; MVR, multivariable regression; PGC, Psychiatric Genomics Consortium; SCZ, Schizophrenia; $P_{thr}$, *P*-value threshold; woADHD; without ADHD

Sets of independent genetic variants were selected from ASD(iPSYCH, woADHD) at different *P*-value thresholds ($P_{thr}$<0.0015, $P_{thr}$<0.05). In ASD-MVR models (Figure 1a), cross-disorder effects were estimated with MDD, SCZ or BD SNP estimates. SNP estimates were extracted from ASD(iPSYCH, woADHD), MDD(PGC), SCZ(PGC), BD(PGC) and EA(SSGAC) GWAS statistics. ASD-specific MVR effects quantify the change in years-of-schooling per log-odds ASD conditional on cross-disorder effects. Latter reflect the change in years-of-schooling per log-odds liability in either MDD, SCZ or BD, as captured by ASD variant sets. For ASD-MVR, SNP estimates were aligned according to ASD risk. The multiple testing threshold is *P*<0.0042. The model fit of MVRs and univariable regressions was compared with likelihood-ratio tests.

Supplementary Table 17: Follow-up ADHD-MVR ($P_{thr}$<0.0015; $P_{thr}$<0.05), analyses with MDD, SCZ and BD SNP estimates (predictor)

| Variant selection | | | Cross-disorder | Intercept | | ADHD-specific effect | | Cross-disorder effect | | VIF | Model fit compared to a single model | |
|---|---|---|---|---|---|---|---|---|---|---|---|---|
| Variants | $N_{SNPs}$ | $P_{thr}$ | | $\beta_{int}$(SE) | *P* | MVR $\beta_{ADHD}$(SE) | *P* | MVR $\beta_{\otimes}$(SE) | *P* | | $\Delta R^2$ (%) | $\Delta$deviance, *P* |
| ADHD (iPSYCH) | 2,567 | 0.0015 | MDD (PGC) | -0.001(2x10$^{-4}$) | <1x10$^{-10}$ | -0.005(0.003) | 0.11 | NA | | NA | NA | |
| | | | | -0.001(2x10$^{-4}$) | 4x10$^{-10}$ | -0.002(0.003) | 0.56 | $\beta_{\otimes MDD}$=-0.033(0.007) | 5x10$^{-7}$ | 1.04 | 0.98 | 76.87, 4x10$^{-7}$ |
| | 38,093 | 0.05 | | -0.001(4x10$^{-5}$) | <1x10$^{-10}$ | -0.005(0.001) | 1x10$^{-10}$ | NA | | NA | NA | |
| | | | | -0.001(4x10$^{-5}$) | <1x10$^{-10}$ | -0.004(0.001) | 5x10$^{-7}$ | $\beta_{\otimes MDD}$=-0.013(0.001) | <1x10$^{-10}$ | 1.02 | 0.23 | 181.10, <1x10$^{-10}$ |
| | 2,278 | 0.0015 | SCZ (PGC) | -0.001(3x10$^{-4}$) | 7x10$^{-7}$ | -0.007(0.004) | 0.056 | NA | | NA | NA | |
| | | | | -0.001(3x10$^{-4}$) | 9x10$^{-7}$ | -0.007(0.004) | 0.059 | $\beta_{\otimes SCZ}$=-0.006(0.004) | 0.14 | 1.00 | 0.09 | 6.91, 0.14 |
| | 31,395 | 0.05 | | -0.001(5x10$^{-5}$) | <1x10$^{-10}$ | -0.007(0.001) | <1x10$^{-10}$ | NA | | NA | NA | |
| | | | | -0.001(5x10$^{-5}$) | <1x10$^{-10}$ | -0.007(0.001) | <1x10$^{-10}$ | $\beta_{\otimes SCZ}$=0.003(0.001) | 0.007 | 1.00 | 0.02 | 16.21, 0.007 |
| | 2,340 | 0.0015 | BD (PGC) | -0.001(2x10$^{-4}$) | 1x10$^{-7}$ | -0.007(0.003) | 0.040 | NA | | NA | NA | |
| | | | | -0.001(2x10$^{-4}$) | 8x10$^{-8}$ | -0.007(0.003) | 0.042 | $\beta_{\otimes BD}$=0.007(0.004) | 0.046 | 1.00 | 0.17 | 12.56, 0.046 |
| | 32,788 | 0.05 | | -0.001(5x10$^{-5}$) | <1x10$^{-10}$ | -0.007(0.001) | <1x10$^{-10}$ | NA | | NA | NA | |
| | | | | -0.001(5x10$^{-5}$) | <1x10$^{-10}$ | -0.007(0.001) | <1x10$^{-10}$ | $\beta_{\otimes BD}$=0.008(0.001) | <1x10$^{-10}$ | 1.00 | 0.29 | 205.72, <1x10$^{-10}$ |

Abbreviations: ADHD, Attention-Deficit/Hyperactivity Disorder; BD, Bipolar Disorder; iPSYCH, The Lundbeck Foundation Initiative for Integrative Psychiatric Research; MDD; Major Depressive Disorder; MVR, multivariable regression; PGC, Psychiatric Genomics Consortium; SCZ, Schizophrenia; $P_{thr}$, *P*-value threshold; woADHD; without ADHD

Sets of independent genetic variants were selected from ADHD(iPSYCH) GWAS statistics at different *P*-value thresholds ($P_{thr}$<0.0015, $P_{thr}$<0.05). In ADHD-MVR models (Figure 1b) cross-disorder effects were estimated with MDD, SCZ or BD SNP estimates. SNP estimates were extracted from ADHD(iPSYCH), MDD(PGC), SCZ(PGC), BD(PGC) and EA(SSGAC) GWAS statistics. ADHD-specific MVR effects quantify the change in years-of-schooling per log-odds ADHD liability conditional on cross-disorder effects. Latter reflect the change in years-of-schooling per log-odds liability in either MDD, SCZ or BD, as captured by ADHD variant sets. For ADHD-MVR, SNP estimates were aligned according to ADHD risk. The multiple testing threshold is *P*<0.0042. The model fit of MVRs and univariable regressions was compared with likelihood-ratio tests.

Supplementary Table 18: Simulation of a hypothetical multifactorial model of EA, ASD and ADHD interrelationships

| Parameter | | Simulated parameter | Estimated parameter (SE) |
|---|---|---|---|
| SNP-$h^2$ | EA | 0.25 | 0.34 (0.06) |
| | ADHD liability | 0.50 | 0.42 (0.06) |
| | ASD liability | 0.50 | 0.40 (0.06) |
| $e^2$ | EA | 0.75 | 0.66 (0.06) |
| | ADHD liability | 0.50 | 0.58 (0.06) |
| | ASD liability | 0.50 | 0.60 (0.06) |
| Genetic factor loadings | $a_{11}$ | 0.50 | -0.58 (0.05) |
| | $a_{21}$ | -0.35 | 0.30 (0.07) |
| | $a_{31}$ | 0.20 | -0.14 (0.07) |
| | $a_{22}$ | 0.62 | -0.58 (0.05) |
| | $a_{32}$ | 0.14 | -0.11 (0.08) |
| | $a_{33}$ | 0.66 | -0.61 (0.05) |
| Residual factor loadings | $e_{11}$ | 0.87 | -0.81 (0.04) |
| | $e_{21}$ | 0.00 | 0.01 (0.05) |
| | $e_{31}$ | 0.00 | -0.03 (0.05) |
| | $e_{22}$ | 0.71 | 0.76 (0.04) |
| | $e_{32}$ | 0.00 | 0.00 (0.05) |
| | $e_{33}$ | 0.71 | -0.77 (0.04) |
| $r_g$ | EA, ADHD | -0.49 | -0.46 (0.11) |
| | EA, ASD | 0.28 | 0.23 (0.11) |
| | ADHD, ASD | 0.04 | 0.04 (0.10) |

Abbreviations: ADHD, Attention-Deficit/Hyperactivity Disorder; ASD, Autism Spectrum Disorder; EA, educational attainment; SNP-$h^2$, Single Nucleotide Polymorphism heritability; $e^2$, residual variance; $r_g$, genetic correlation

We simulated three continuous interrelated measures corresponding to EA, ADHD liability and ASD liability (N=6000 each), informed by unconstrained LDSC genetic correlations ($r_g$) using GWAS statistics for EA, ADHD(iPSYCH) and ASD(PGC). Residual correlations were assumed to be absent, given the independence of the respective GWAS statistics. Note that simulated SNP-$h^2$ estimates were increased, compared to the observed values, to reduce the computational burden.

Supplementary Figures

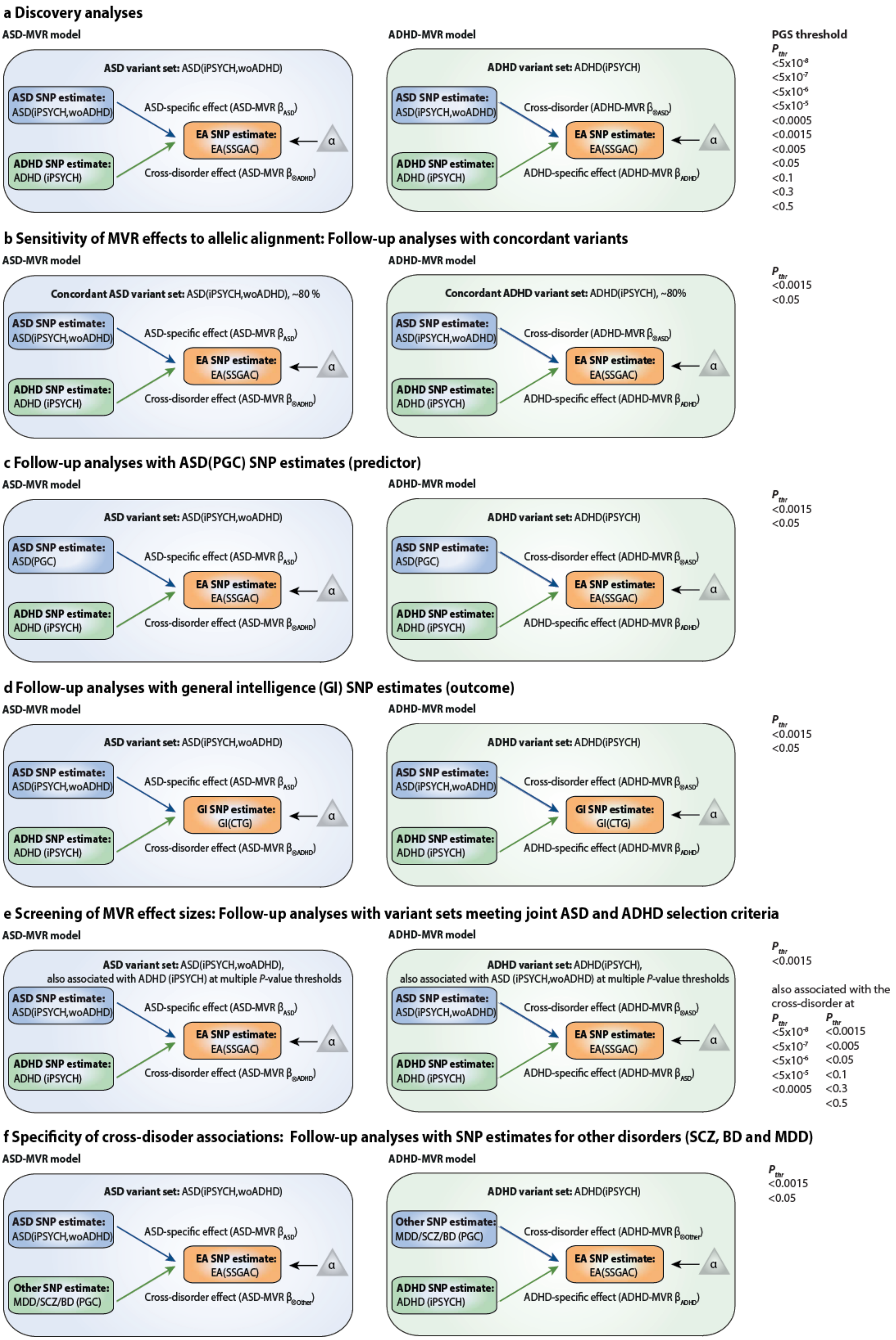
a Discovery analyses
ASD-MVR model
ASD variant set: ASD(iPSYCH,woADHD)
ASD SNP estimate: ASD(iPSYCH,woADHD)
ASD-specific effect (ASD-MVR βASD)
EA SNP estimate: EA(SSGAC)
α
ADHD SNP estimate: ADHD (iPSYCH)
Cross-disorder effect (ASD-MVR β⊗ADHD)
ADHD-MVR model
ADHD variant set: ADHD(iPSYCH)
ASD SNP estimate: ASD(iPSYCH,woADHD)
Cross-disorder (ADHD-MVR β⊗ASD)
EA SNP estimate: EA(SSGAC)
α
ADHD SNP estimate: ADHD (iPSYCH)
ADHD-specific effect (ADHD-MVR βADHD)
PGS threshold
Pthr
<5x10^-8
<5x10^-7
<5x10^-6
<5x10^-5
<0.0005
<0.0015
<0.005
<0.05
<0.1
<0.3
<0.5
b Sensitivity of MVR effects to allelic alignment: Follow-up analyses with concordant variants
ASD-MVR model
Concordant ASD variant set: ASD(iPSYCH,woADHD), ~80 %
ASD SNP estimate: ASD(iPSYCH,woADHD)
ASD-specific effect (ASD-MVR βASD)
EA SNP estimate: EA(SSGAC)
α
ADHD SNP estimate: ADHD (iPSYCH)
Cross-disorder effect (ASD-MVR β⊗ADHD)
ADHD-MVR model
Concordant ADHD variant set: ADHD(iPSYCH), ~80%
ASD SNP estimate: ASD(iPSYCH,woADHD)
Cross-disorder (ADHD-MVR β⊗ASD)
EA SNP estimate: EA(SSGAC)
α
ADHD SNP estimate: ADHD (iPSYCH)
ADHD-specific effect (ADHD-MVR βADHD)
Pthr
<0.0015
<0.05
c Follow-up analyses with ASD(PGC) SNP estimates (predictor)
ASD-MVR model
ASD variant set: ASD(iPSYCH,woADHD)
ASD SNP estimate: ASD(PGC)
ASD-specific effect (ASD-MVR βASD)
EA SNP estimate: EA(SSGAC)
α
ADHD SNP estimate: ADHD (iPSYCH)
Cross-disorder effect (ASD-MVR β⊗ADHD)
ADHD-MVR model
ADHD variant set: ADHD(iPSYCH)
ASD SNP estimate: ASD(PGC)
Cross-disorder effect (ADHD-MVR β⊗ASD)
EA SNP estimate: EA(SSGAC)
α
ADHD SNP estimate: ADHD (iPSYCH)
ADHD-specific effect (ADHD-MVR βADHD)
Pthr
<0.0015
<0.05
d Follow-up analyses with general intelligence (GI) SNP estimates (outcome)
ASD-MVR model
ASD variant set: ASD(iPSYCH,woADHD)
ASD SNP estimate: ASD(iPSYCH,woADHD)
ASD-specific effect (ASD-MVR βASD)
GI SNP estimate: GI(CTG)
α
ADHD SNP estimate: ADHD (iPSYCH)
Cross-disorder effect (ASD-MVR β⊗ADHD)
ADHD-MVR model
ADHD variant set: ADHD(iPSYCH)
ASD SNP estimate: ASD(iPSYCH,woADHD)
Cross-disorder effect (ADHD-MVR β⊗ASD)
GI SNP estimate: GI(CTG)
α
ADHD SNP estimate: ADHD (iPSYCH)
ADHD-specific effect (ADHD-MVR βADHD)
Pthr
<0.0015
<0.05
e Screening of MVR effect sizes: Follow-up analyses with variant sets meeting joint ASD and ADHD selection criteria
ASD-MVR model
ASD variant set: ASD(iPSYCH,woADHD),
also associated with ADHD (iPSYCH) at multiple P-value thresholds
ASD SNP estimate: ASD(iPSYCH,woADHD)
ASD-specific effect (ASD-MVR βASD)
EA SNP estimate: EA(SSGAC)
α
ADHD SNP estimate: ADHD (iPSYCH)
Cross-disorder effect (ASD-MVR β⊗ADHD)
ADHD-MVR model
ADHD variant set: ADHD(iPSYCH),
also associated with ASD (iPSYCH,woADHD) at multiple P-value thresholds
ASD SNP estimate: ASD(iPSYCH,woADHD)
Cross-disorder effect (ADHD-MVR β⊗ASD)
EA SNP estimate: EA(SSGAC)
α
ADHD SNP estimate: ADHD (iPSYCH)
ADHD-specific effect (ADHD-MVR βASD)
Pthr
<0.0015
also associated with the cross-disorder at
Pthr Pthr
<5x10^-8 <0.0015
<5x10^-7 <0.005
<5x10^-6 <0.05
<5x10^-5 <0.1
<0.0005 <0.3
<0.5
f Specificity of cross-disoder associations: Follow-up analyses with SNP estimates for other disorders (SCZ, BD and MDD)
ASD-MVR model
ASD variant set: ASD(iPSYCH,woADHD)
ASD SNP estimate: ASD(iPSYCH,woADHD)
ASD-specific effect (ASD-MVR βASD)
EA SNP estimate: EA(SSGAC)
α
Other SNP estimate: MDD/SCZ/BD (PGC)
Cross-disorder effect (ASD-MVR β⊗Other)
ADHD-MVR model
ADHD variant set: ADHD(iPSYCH)
Other SNP estimate: MDD/SCZ/BD (PGC)
Cross-disorder effect (ADHD-MVR β⊗Other)
EA SNP estimate: EA(SSGAC)
α
ADHD SNP estimate: ADHD (iPSYCH)
ADHD-specific effect (ADHD-MVR βADHD)
Pthr
<0.0015
<0.05

### Supplementary Figure 1: Study design of interlinked MVR analyses

In principle, we applied a weighted multivariable regression and regressed SNP estimates for EA (dependent variable) jointly on SNP estimates for ASD (independent variable) and SNP estimates for ADHD (independent variable), including a regression intercept (α). For all MVR analyses, ASD-related and ADHD-related variant sets were selected from ASD(iPSYCH,woADHD) and ADHD(iPSYCH) GWAS statistics, respectively, using multiple *P*-value thresholds ($5x10^{-8} \leq P_{thr} < 0.5$). For the discovery analyses (a), this involved a screening with multiple variant sets across a range of 11 *P*-value thresholds ($P_{thr}$, $5x10^{-8}$; $5x10^{-7}$; $5x10^{-6}$; $5x10^{-5}$; 0.0005; 0.0015; 0.005; 0.05; 0.1; 0.3; 0.5) for each neurodevelopmental disorder. We predominantly focus on two *P*-value thresholds: (i) $P_{thr}<0.0015$, consistent with guidelines for validating genetic instrument strength (F-statistic<10)[4] and conservative selection thresholds recommended for related polygenic scoring approaches[5], and (ii) $P_{thr}<0.05$, a less stringent threshold with the aim to increase the statistical power and precision of MVR estimates. All variants were restricted to common (minor allele frequency>0.01), independent (linkage disequilibrium-$r^2<0.25$ within ±500 kb[6]) and well-imputed (Imputation quality(INFO)[7]>0.7) SNPs. For MVR models (a,b,e), SNP estimates were extracted from ASD(iPSYCH, woADHD), ADHD(iPSYCH) and EA(SSGAC) GWAS statistics, for MVR models (c) from ASD(PGC), ADHD(iPSYCH) and EA(SSGAC), for MVR models (d) from ASD(PGC), ADHD(iPSYCH) and GI(CTG) and for MVR models (f) from ASD(iPSYCH, woADHD), ADHD(iPSYCH), MDD(PGC), SCZ(PGC), BP(PGC) and EA(SSGAC).

Abbreviations: ADHD, Attention-Deficit/Hyperactivity Disorder; ASD, Autism Spectrum Disorder; BD, Bipolar Disorder; GWAS, genome-wide association study, iPSYCH, The Lundbeck Foundation Initiative for Integrative Psychiatric Research; MDD; Major Depressive Disorder; MVR, multivariable regression; $P_{thr}$, *P*-value threshold; PGC, Psychiatric Genomics Consortium; SCZ, Schizophrenia; SSGAC, Social Science Genetic Consortium; woADHD; without ADHD

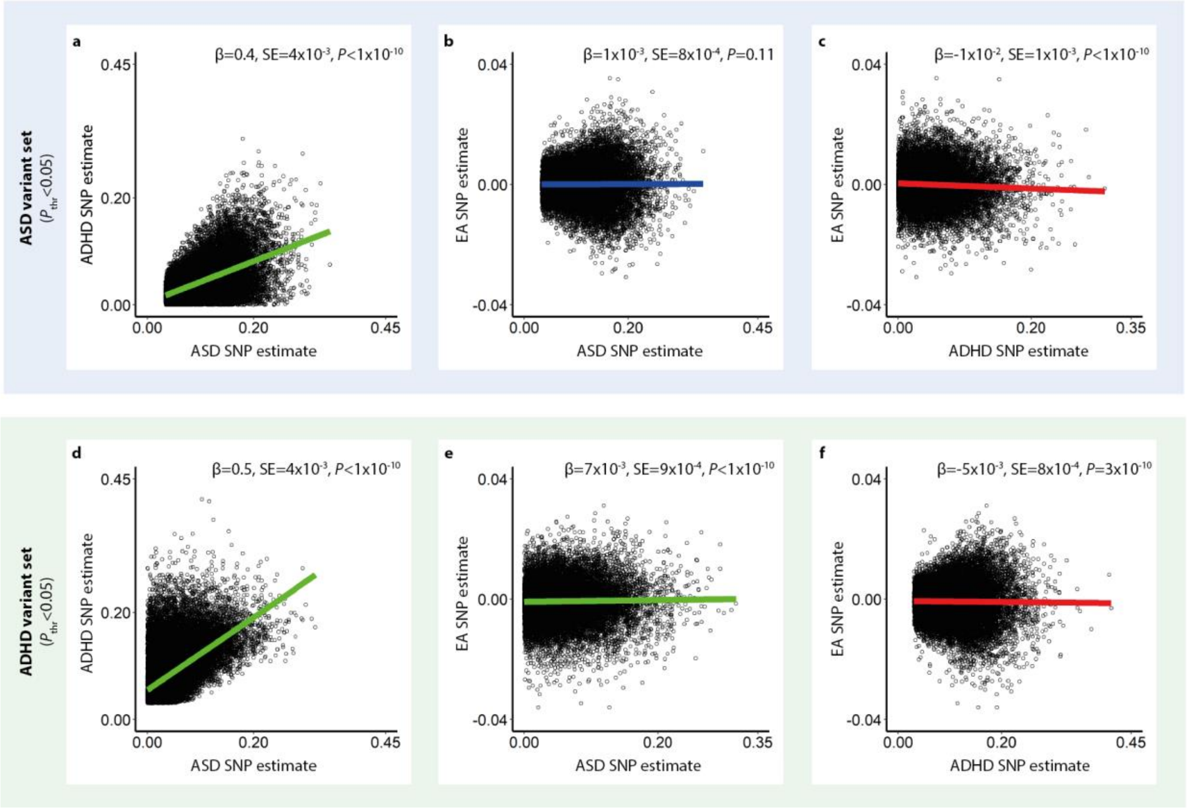

Supplementary Figure 2: Bivariate analyses of ASD, ADHD and EA genetic effects using ASD and ADHD concordant variant sets

Sets of independent ASD and ADHD genetic variants passing a *P*-value threshold of 0.05 were selected from ASD(iPSYCH, woADHD) and ADHD(iPSYCH) GWAS statistics, respectively. Corresponding ASD, ADHD and EA SNP estimates were subsequently extracted from ASD(iPSYCH, woADHD) and ADHD(iPSYCH) and EA(SSGAC) GWAS statistics. Only genetic variants with the same risk allele for both ASD and ADHD were included. Thus, SNP estimates are aligned to increase risk for both disorders. Using single weighted regression models and ASD variants (a-c), SNP estimates for ADHD were regressed on SNP estimates for ASD (a), SNP estimates for EA were regressed on SNP estimates for ASD (b) and SNP estimates for EA were regressed on SNP estimates for ADHD (c). Using single weighted regression models and ADHD variants (d-f), SNP estimates for ADHD were regressed on SNP estimates for ASD (d), SNP estimates for EA were regressed on SNP estimates for ASD (e) and SNP estimates for EA were regressed on SNP estimates for ADHD (f). All regressions allowed for an intercept. A green regression line denotes a positive relationship between SNP estimates, a blue regression line reflects no significant relationship and a red line indicates a negative relationship. Regression beta estimates, corresponding standard errors (SE) and *P*-values are shown for each regression model.

Abbreviations: ASD, Autism Spectrum Disorder; ADHD, Attention-Deficit/Hyperactivity Disorder; EA, educational attainment; GWAS, genome-wide association study, $P_{thr}$, *P*-value threshold

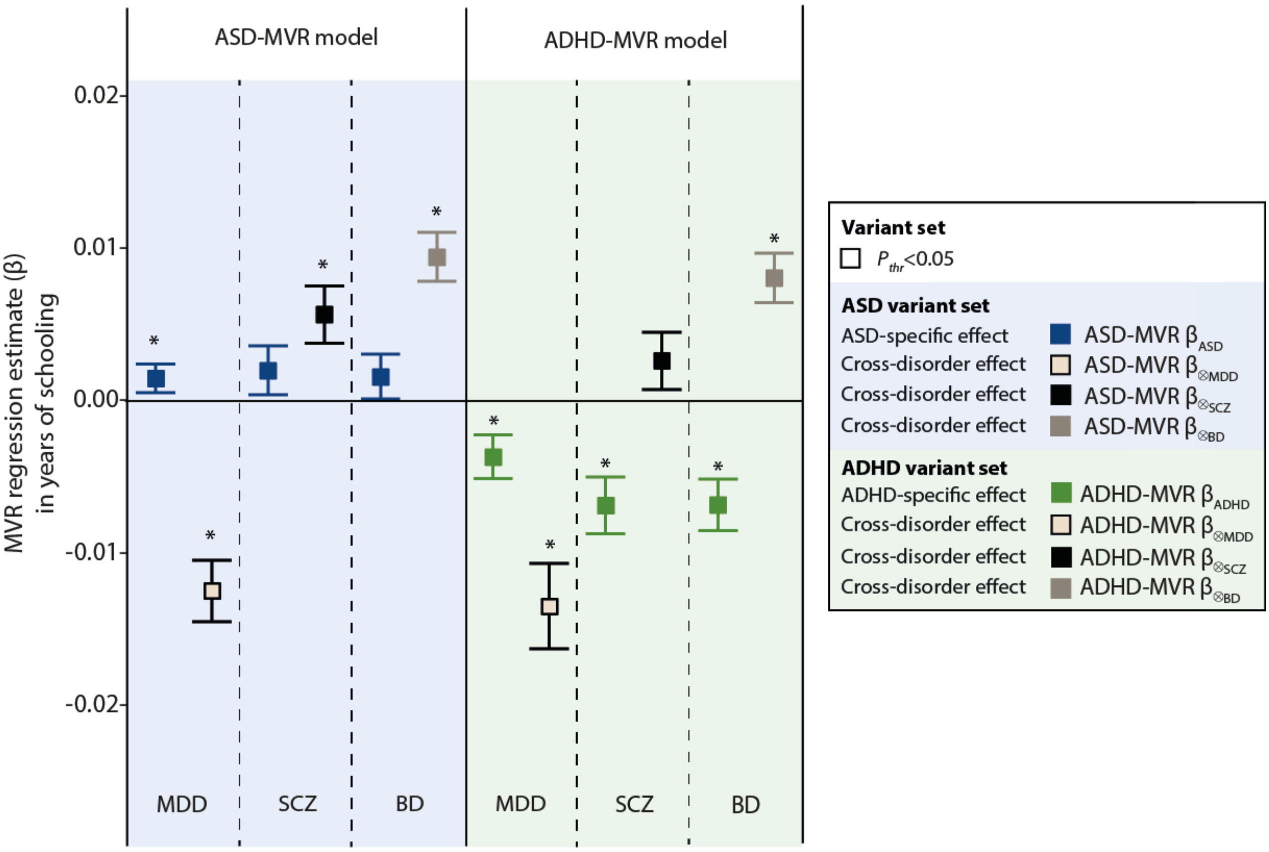

Supplementary Figure 3: Specificity of cross-disorder associations with educational attainment: Follow up analyses with MDD, SCZ and BD SNP estimates (predictor)

Sets of independent ASD and ADHD genetic variants were selected from ASD(iPSYCH, woADHD) and ADHD(iPSYCH) summary statistics, respectively, at $P_{thr}$<0.05 (MVR effects at $P_{thr}$<0.0015 were omitted for clarity). Corresponding SNP estimates for ASD, ADHD and EA were subsequently extracted from ASD(iPSYCH, woADHD), ADHD(iPSYCH) and EA(SSGAC) GWAS statistics, respectively, in addition to MDD, SCZ and BD SNP estimates (PGC summary statistics). For ASD-MVR models (using ASD variant sets), ASD-specific effects (ASD-MVR βASD) are shown in addition to unspecific cross-disorder associations shared with MDD (ASD-MVR β⨂MDD), SCZ (ASD-MVR β⨂SCZ) and BD (ASD-MVR β⨂MDD). For ADHD-MVR models (using ADHD variant sets), ADHD-specific effects (ADHD-MVR βADHD) are shown in addition to unspecific cross-disorder associations shared with MDD (ADHD-MVR β⨂MDD), SCZ (ADHD-MVR β⨂SCZ) and BD (ADHD-MVR β⨂MDD). For, ASD-MVR, SNP effects were aligned according to ASD risk, for ADHD-MVR SNP effects according to ADHD risk. All MVR effects are presented as changes in years-of-schooling per increase in log-odds liability for psychiatric disorder. Bars represent 95% confidence intervals.

*MVR effects passing the multiple testing threshold of P<0.0042.

Abbreviations: ADHD, Attention-Deficit/Hyperactivity Disorder; ASD, Autism Spectrum Disorder; BD, Bipolar Disorder; MDD; Major Depressive Disorder; MVR, multivariable regression; SCZ, Schizophrenia; $P_{thr}$, P-value threshold.

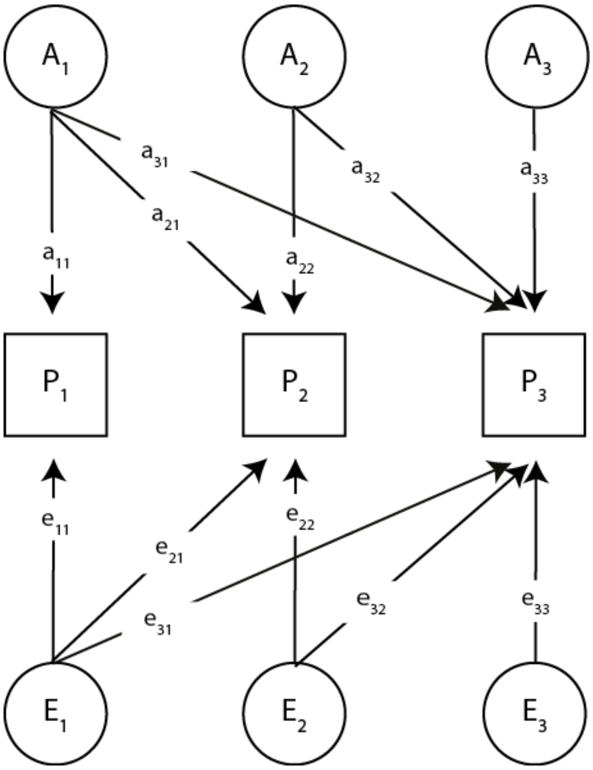

Supplementary Figure 4: Path diagram for a trivariate trait

The variance/covariance structure of multivariate trait consisting of three standardised measures P1, P2 and P3 can be described using a Cholesky decomposition consisting of three genetic factors (A1, A2 and A3) and three residual factors (E1, E2 and E3), shown here with genetic and residual factor loadings

The observed phenotypic measures are represented by squares, while all latent genetic and residual factors are represented by a circle. Single headed arrows ('paths') denote causal relationships between variables and are shown for genetic factor loadings (a) and residual factor loadings (e). Note that the variance of latent variables is constrained to unit variance, this is omitted from the diagrams to improve clarity.

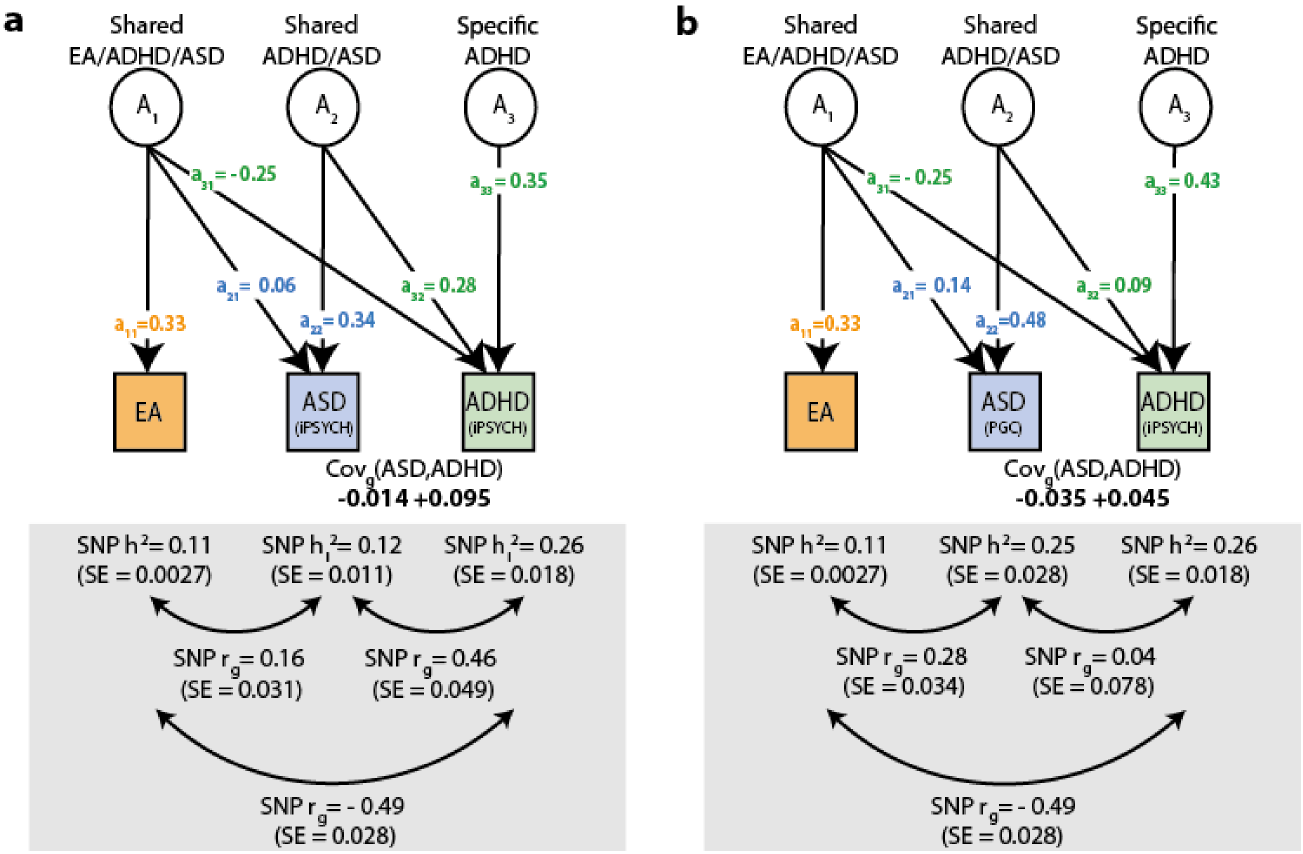

### Supplementary Figure 5: Multi-factor model of genetic interrelations between ASD, ADHD and educational attainment (allowing for ADHD-specific genetic influences)

The model predicts two sources of shared genetic influences between ASD and ADHD, as captured by common variants within an infinitely large population. The first genetic factor (A1, shared EA/ADHD/ASD) refers to shared genetic variation between EA, ADHD and ASD. It allows for a negative genetic covariance between ASD and ADHD. The second genetic factor (A2, shared ADHD/ASD) acts independently of A1, explaining positive genetic covariance between ASD and ADHD.

Each factor loading ("a") for the Cholesky decomposition of a trivariate trait is described in the Methods. **(a)** Multi-factor model consistent with ASD(iPSYCH), ADHD(iPSYCH) and EA(SSGAC) summary statistics. **(b)** Multi-factor model consistent with ASD(PGC), ADHD(iPSYCH) and EA(SSGAC) summary statistics.

Factor loadings ("a") were derived from LDSC SNP-heritability and genetic correlations (grey boxes). Shared ADHD/ASD genetic influences (A2) were modelled allowing for ADHD-specific effects (A3). Phenotypic measures are represented by squares, while latent genetic factors are represented by circles. Single headed arrows denote genetic factor loadings ("a"), double-headed arrows genetic correlations ("$r_g$"). Residual influences and unit variances for latent variables were omitted.

Abbreviations: EA, educational attainment; ADHD, Attention-Deficit/Hyperactivity Disorder; ASD, Autism Spectrum Disorder; iPSYCH, The Lundbeck Foundation Initiative for Integrative Psychiatric Research; PGC, Psychiatric Genomics Consortium; SNP $h^2$, SNP heritability; SNP $r_g$, SNP genetic correlation, $cov_g$, genetic covariance

### References

1. Savage JE, Jansen PR, Stringer S, et al. Genome-wide association meta-analysis in 269,867 individuals identifies new genetic and functional links to intelligence. *Nature Genetics*. 2018;50(7):912-919. doi:10.1038/s41588-018-0152-6

2. Bulik-Sullivan BK, Loh P-R, Finucane HK, et al. LD Score regression distinguishes confounding from polygenicity in genome-wide association studies. *Nat Genet*. 2015;47(3):291-295. doi:10.1038/ng.3211

3. Bulik-Sullivan B, Finucane HK, Anttila V, et al. An atlas of genetic correlations across human diseases and traits. *Nat Genet*. 2015;47:1236–1241. doi:10.1038/ng.3406

4. Pierce BL, Ahsan H, Vanderweele TJ. Power and instrument strength requirements for Mendelian randomization studies using multiple genetic variants. *Int J Epidemiol*. 2011;40(3):740-752. doi:10.1093/ije/dyq151

5. Wray NR, Lee SH, Mehta D, Vinkhuyzen AAE, Dudbridge F, Middeldorp CM. Research Review: Polygenic methods and their application to psychiatric traits. *J Child Psychol Psychiatr*. 2014;55(10):1068-1087. doi:10.1111/jcpp.12295

6. Purcell S, Neale B, Todd-Brown K, et al. PLINK: A Tool Set for Whole-Genome Association and Population-Based Linkage Analyses. *Am J Hum Genet*. 2007;81(3):559-575.

7. Howie BN, Donnelly P, Marchini J. A Flexible and Accurate Genotype Imputation Method for the Next Generation of Genome-Wide Association Studies. *PLOS Genetics*. 2009;5(6):e1000529. doi:10.1371/journal.pgen.1000529
